# DNMT1 Coordinates PV Interneuron–Glia Coupling to Maintain Cortical Network Stability and Regulate Behavior

**DOI:** 10.1101/2025.11.18.689032

**Authors:** Jenice Linde, Can Bora Yildiz, Katharina Vöhringer, Gerion Nabbefeld, Severin Graff, Daniel Pensold, Julia Reichard, Marie Hermanns, Christoph Weber-Hamacher, Marc Spehr, Björn Kampa, Anja Urbach, Simon Musall, Geraldine Zimmer-Bensch

**Author notes:** Equal contribution (Jenice Linde, Can Bora Yildiz). Equal contribution (Simon Musall, Geraldine Zimmer-Bensch).

## Abstract

Parvalbumin (PV) interneurons are central to cortical network stability and psychiatric vulnerability. Here, we identify DNA methyltransferase 1 (DNMT1) as a key epigenetic regulator linking PV interneuron function to glial and extracellular matrix remodeling. Conditional PV-specific *Dnmt1* deletion combined with single-cell RNA-seq, in vivo electrophysiology, histology, and behavioral analyses revealed that loss of DNMT1 increases PV spiking activity but reduces inhibitory efficacy, leading to network desynchronization and depression- and anxiety-like behavior in mice. These physiological alterations were accompanied by broad, non-cell-autonomous transcriptional changes in astrocytes and oligodendroglial populations, prominently affecting pathways involved in perineuronal-net (PNN) organization and neuron-glia communication. Cell-cell interaction analyses revealed disrupted NRXN-NLGN, TNR-integrin, and semaphoring signaling, consistent with weakened perisomatic adhesion and PNN integrity. Together, our findings demonstrate that DNMT1 maintains inhibitory circuit stability through cell-autonomous regulation of PV interneuron function, which secondarily shapes glial transcriptional states and extracellular scaffolds to preserve cortical network synchronization and emotional behavior.

## Introduction

GABAergic interneurons are critical regulators of cortical circuit dynamics, ensuring precise control over pyramidal neuron activity and maintaining the excitation/inhibition (E/I) balance essential for healthy brain function. Disruption of inhibitory signaling contribute to maladaptive network states and have been implicated in a range of neuropsychiatric disorders, including schizophrenia and major depressive disorder (MDD)^1–3^. However, the etiology of such disorders is inherently multifactorial, due to interactions between genetic predispositions, environmental stressors, and epigenetic mechanisms that converge to shape disease vulnerability^3^. The exact nature of GABAergic dysfunction in MDD therefore remains debated, with contradictory findings further complicating the establishment of a unified mechanistic framework^4,5^.

Epigenetic modifications, such as DNA methylation mediated by DNA methyltransferases (DNMTs), form a crucial interface between external stimuli and neuronal function. Aberrant DNA methylation has been documented across multiple neuronal subtypes in psychiatric and neurological conditions, including schizophrenia, epilepsy, and MDD^6,7^. Among DNMTs, DNMT1 has emerged as a key regulator of inhibitory circuit function: altered DNMT1 expression in cortical interneurons has been linked to schizophrenia and disrupted GABAergic signaling^3^.

Within the diverse population of inhibitory neurons, parvalbumin-positive (PV) interneurons are especially important to regulate network activity. As fast-spiking interneurons, they synchronize network oscillations and gate excitatory inputs, thereby maintaining cortical E/I balance^8^. Alterations in PV interneuron number or function have been causally linked to neuropsychiatric diseases^9,10^, and their vulnerability to chronic stress makes them prime candidates for involvement in MDD pathology^11^. Our previous work also demonstrated that DNMT1 exerts transcriptional control in postmitotic cortical interneurons^12^ and regulates synaptic transmission in PV interneurons by modulating endocytosis-dependent GABA reuptake at presynaptic terminals, thereby fine-tuning inhibitory output^13^.

Beyond their intrinsic properties, PV interneurons are profoundly shaped by interactions with surrounding glial cells and the extracellular matrix. These include astrocytes and oligodendrocytes, as well as extracellular matrix components, such as perineuronal nets (PNNs). PNNs tightly ensheathe PV interneurons, stabilize synaptic contacts, and limit plasticity, thereby influencing both inhibitory efficacy and network stability^14,15^. Disruption of glial support or PNN integrity has therefore been increasingly recognized as an important contributor to PV interneuron dysfunction^16^ and psychiatric disease vulnerability^17^.

In this study, we investigated the cell-autonomous and non-cell-autonomous consequences of DNMT1 function in PV interneurons to reveal how epigenetic dysregulation contributes to cortical circuit instability and disease vulnerability. To this end, we used conditional PV-specific *Dnmt1* knockout mice and combined single-cell transcriptomics, electrophysiology, histological analyses, and behavioral assays. *Dnmt1* deletion reduced the inhibitory impact of PV interneurons on network activity in vivo and altered oligodendroglial transcriptional programs converging on PNN regulation and neuron-glia communication. This dual effect identifies DNMT1 as a central epigenetic coordinator of inhibitory circuit function, linking intrinsic PV interneuron activity with extrinsic glial and extracellular matrix support. Together, our findings reveal how epigenetic perturbations in inhibitory neurons propagate across cellular domains, thereby destabilizing cortical networks and promoting behavioral phenotypes reminiscent of MDD.

## Results

### *Dnmt1* deletion in PV interneurons alters cortical network activity across different cortical areas

Building on a PV-specific conditional *Dnmt1* knockout mouse line^13,18^, we already demonstrated that DNMT1 regulates GABAergic transmission in PV interneurons through regulation of endocytosis-associated gene expression^13^. Given that cortical interneuron dysfunction and dysregulated *Dnmt1* expression have both been implicated in neuropsychiatric diseases^19^, we next examined the functional consequences of *Dnmt1* deletion in PV interneurons at the network level. To this end, we performed in vivo electrophysiological recordings in the primary visual cortex (V1) of awake, behaving *Pvalb-Cre/tdTomato/Dnmt1 loxP^2^* knockout (KO) and *Pvalb-Cre/tdTomato* control mice using Neuropixels probes (Fig. 1a).

**Figure 1:**
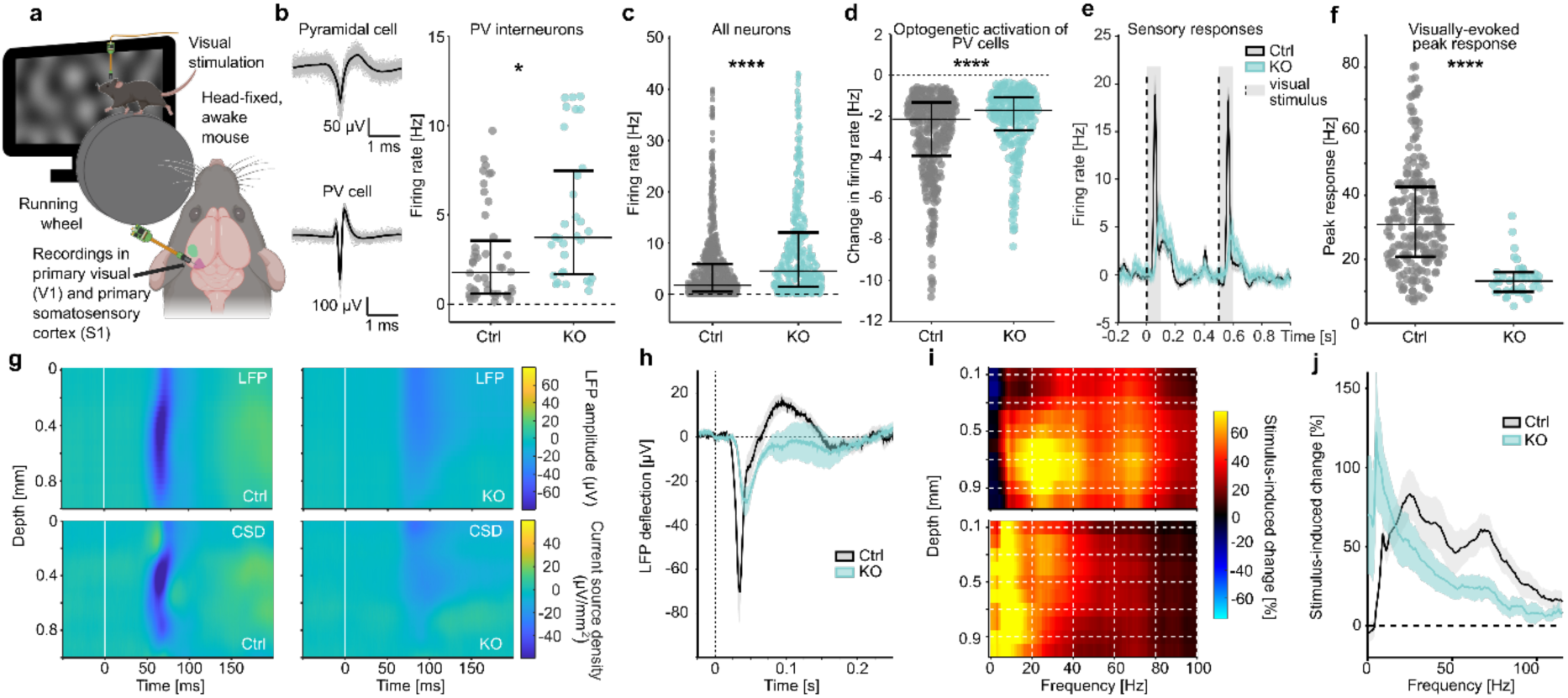
*Dnmt1*-knockout in PV interneurons alters spontaneous and induced neuronal activity. We performed electrophysiological recordings in the primary visual cortex (V1) of awake *Pvalb-Cre/tdTomato* control (Ctrl) and *Pvalb-Cre/tdTomato/Dnmt1 loxP^2^* (KO) mice using Neuropixels probes (*n* = 8 recordings from 2 mice per genotype). **(a)** Schematic illustration of the experimental setup and recording sites. **(b)** Fast-spiking PV interneurons were identified via optotagging (see example spike waveforms, left). Their spontaneous firing rates were significantly increased in KO mice (*n*_Ctrl_ = 54 and *n*_KO_ = 37 neurons; Wilcoxon *rank-sum* test: *p* = 0.0297). **(c)** Spontaneous firing rates of V1 neurons were significantly higher in KO mice compared to control mice (*n*_Ctrl_ = 838 and *n*_KO_ = 448 neurons; Wilcoxon *rank-sum* test: *p* = 5.72×10^-19^). **(d)** Relative change in the firing rate of V1 neurons inhibited by optogenetic activation of PV interneurons. PV-induced inhibition was significantly weaker in KO mice (*n*_Ctrl_=332 and *n*_KO_=238 neurons; Wilcoxon *rank-sum* test: *p* = 3×10^-6^). **(e)** Average firing rates of V1 neurons in response to visual stimulation in control and knockout mice. Visual stimuli were 0.1-s-long, full-field presentations of low-frequency noise patterns, presented at time point 0 and after 0.5 seconds. Shading shows the standard error of means (SEM) across neurons. **(f)** Peak response firing rate of V1 neurons after visual stimulation. KO mice had significantly weaker sensory responses compared to control mice (*n*_Ctrl_ = 293 and *n*_KO_ = 74 neurons; Wilcoxon *rank-sum* test: *p* = 1.1×10^-15^). **(g)** Local field potentials (LFP, upper panels) and current source densities (CSD, lower panels) of control and *Dnmt1* KO mice across the whole depth of the cortex. White lines indicate visual stimulus onset. **(h)** LFPs in V1 in response to visual stimulation at time point 0, averaged across cortical layers. Shading shows SEM across recordings. **(i–j)** KO mice displayed strongly diminished gamma oscillations (30–90 Hz) in comparison to control mice. **(i)** Relative change in spectral power compared to baseline activity. Visually evoked oscillations in control mice matched the gamma band (30–90 Hz), while in KO mice oscillations of much lower frequencies (< 20 Hz) occurred. **(j)** Change in spectral power within the cortex displayed as mean ± SEM. The direct comparison between control and *Dnmt1* KO mice revealed the shift towards lower-frequency oscillations in KO mice. Panels b–f display individual data points with median (centre line) and interquartile range (upper and lower lines). Ctrl = control, CSD = current source density, KO = knockout, LFP = local field potential, PV = parvalbumin, V1 = primary visual cortex; * *p* < 0.5, ** *p* < 0.01, *** *p* < 0.001, **** *p* < 0.0001.

To selectively modulate PV interneuron activity, we used viral injections to induce Cre-dependent expression of the blue light-sensitive opsin Channelrhodopsin-2 (ChR2) in V1 PV interneurons. This allowed us to isolate PV spiking activity within the recorded neural population based on their short-latency responses to 10-ms-long blue light pulses (Fig. 1b). Spontaneous firing rates of PV interneurons in *Pvalb-Cre/Dnmt1* KO mice were elevated relative to controls, suggesting that DNMT1 is involved in restraining PV interneuron excitability and confirming our previous in vitro results^13^.

When considering all recorded cortical neurons, including excitatory cells, spontaneous firing rates were also increased in KO mice (Fig. 1c). Moreover, optogenetic activation of PV interneurons strongly inhibited cortical activity in controls but was markedly less effective in KO mice (Fig. 1d). This indicates that the inhibitory impact of PV interneurons on cortical networks was reduced despite their hyperactivity, potentially due to homeostatic network adaptation.

Visual stimulation further revealed altered cortical responses in KO mice. Compared to controls, stimulus-evoked activity in V1 was weaker, temporally delayed, and less precise, with reduced stimulus specificity across cortical layers (Fig. 1a, e–h, Extended Data Fig. 1a–c). In line with earlier findings^12,20^, visual responses in control mice showed a well-defined laminar organization, with the earliest and strongest responses in layer 4 (0.3 – 0.5 mm) and subsequent activation of the superficial and infragranular layers, as evident in the local field potentials (LFPs; Fig 1g, upper panels) and current source densities (CSD; Fig. 1g, lower panels). In contrast, excitatory responses across all cortical layers were reduced in KO mice and the inhibitory rebound component typically associated with feedback inhibition was absent (Fig. 1h). These alterations underscore an impaired contribution of PV interneurons to cortical information processing in KO mice. Consistent with the weakened sensory responses in KO mice (Fig. 1e), we also found fewer stimulus-responsive neurons with a lower response specificity (Extended Data Fig. 1c).

To assess whether these impairments were specific to the visual cortex or generalize to other sensory modalities, we also recorded cortical activity in the primary somatosensory cortex (S1) during whisker stimulation. In controls, tactile input evoked robust and temporally precise responses, peaking in layer 5, followed by a characteristic inhibitory rebound reflecting feedback inhibition (Extended Data Fig. 1d, e). In contrast, *Dnmt1* KO mice showed markedly weaker and less temporally precise responses that were prolonged and lacked the inhibitory rebound that is typically mediated by PV interneurons (Extended Data Fig. 1d, e). Optogenetic activation of PV interneurons during tactile stimulation further confirmed their reduced efficacy: while PV activation strongly suppressed peak responses in controls, it had little effect in KO mice (Extended Data Fig. 1f). These findings therefore match our observations in V1 and indicate that DNMT1 loss in PV interneurons impairs their inhibitory contribution across primary cortical areas.

Next, we extended our recordings to examine higher-order cortical dynamics, focusing on the anterolateral motor cortex (ALM), a prefrontal region engaged in decision-making and evidence accumulation. Because ALM is strongly recruited during goal-directed behaviors, we sought to determine whether DNMT1 loss in PV interneurons also affects higher-order cortical dynamics during task engagement. We therefore trained mice on a visual evidence accumulation task, in which they reported the side with the higher number of cues by licking a corresponding waterspout (Extended Data Fig. 2e). To exclude general learning deficits, we first examined spatial learning and memory by conducting a Morris water maze test. KO and control mice located the hidden platform with similar accuracy and efficiency, showing no genotype-dependent behavioral impairments (Extended Data Fig. 2a–d). Similarly, KO mice acquired the visual evidence accumulation task as efficiently as controls, with no differences in learning curves or performance (Extended Data Fig. 2f, g).

Electrophysiological recordings nevertheless revealed clear alterations in network activity. Similar to V1, ALM neurons in KO mice showed attenuated neural responses to visual stimulation (Extended Data Fig. 2h) and optogenetic activation of PV interneurons again confirmed their reduced inhibitory efficacy (Extended Data Fig. 2i). Importantly, optogenetic activation of PV interneurons impaired decision-making in both genotypes, demonstrating that ALM function remained essential for task performance (Extended Data Fig. 2j).

Together, these findings demonstrate that DNMT1 loss in PV interneurons reduces their inhibitory efficacy across cortical regions, spanning from primary sensory areas to prefrontal circuits critical for decision-making. However, despite these robust physiological alterations, basic perceptual, learning, and memory capacities were preserved, suggesting that compensatory network mechanisms can buffer the functional consequences of impaired PV interneuron control.

### DNMT1 deficiency in PV interneurons impairs gamma synchrony and induces depression-like behavior

PV interneurons are critically involved in the generation of cortical gamma oscillations^21,22^. Strikingly, alongside the altered firing patterns in V1 (Fig. 1g, h) *Dnmt1* KO mice also displayed severely reduced visually evoked gamma oscillations (Fig. 1i, j). Because gamma synchrony depends on the precise timing of PV interneuron output, our findings suggest that DNMT1 is also required to maintain the rhythmic inhibitory drive underlying fast cortical coordination. Importantly, impaired gamma oscillations are a well-documented hallmark of neuropsychiatric disorders, including schizophrenia, bipolar disorder, and major depressive disorder^23^. In patients with depression, reduced gamma power has been associated with cognitive inflexibility, impaired attention, and diminished affective regulation, while animal models consistently link PV dysfunction and gamma desynchronization to depression-like behaviors^22,24,25^.

Notably, bulk RNA sequencing also revealed transcriptional changes in *Pvalb-Cre*/*Dnmt1 KO* mice resembling those in models of depression and anxiety (Extended Data Tab. 1, 2). This raised the question whether DNMT1 loss in PV interneurons might trigger mood disorder-related behaviors. Spontaneous home-cage activity provides a sensitive measure of innate motivation and has been widely used to detect depression- and apathy-like phenotypes in rodents^26^. Monitoring home-cage behavior revealed consistently reduced movements in KO mice (Fig. 2a, b), a change not explained by visual deficits (Extended Data Fig. 2) or motor deficits. KO mice performed normally in rotarod (Fig. 2c) and ladder rung tests^18^, but showed a shorter latency to fall in the wire hang test (Fig. 2d), which coincided with significantly increased body weight (Fig. 2e). Increased body weight is often comorbid with depressive symptoms^27^, and the weight gain of KO mice could therefore reflect reduced physical activity and energy expenditure. In support of this, food intake was unchanged (Extended Data Fig. 3a–c), and appetite-related hormone levels, including ghrelin, insulin, and PYY, were unaffected (Extended Data Fig. 3d). We also detected elevated leptin levels likely reflecting increased adipose tissue rather than altered endocrine regulation^28,29^. Moreover, immunostaining of the arcuate nucleus confirmed that PV-positive cells are largely distinct from the major neuronal populations regulating feeding, showing no overlap with POMC neurons and only partial overlap with AgRP neurons (Extended Data Fig. 4, 5)^30^. This makes it unlikely that *Dnmt1* deletion in PV cells directly interferes with hypothalamic feeding circuits as a driver of weight gain. Together, the obesity and diminished motor activity in KO mice suggest lower energy expenditure and diminished innate motivation consistent with an apathy-like state^31^.

**Figure 2:**
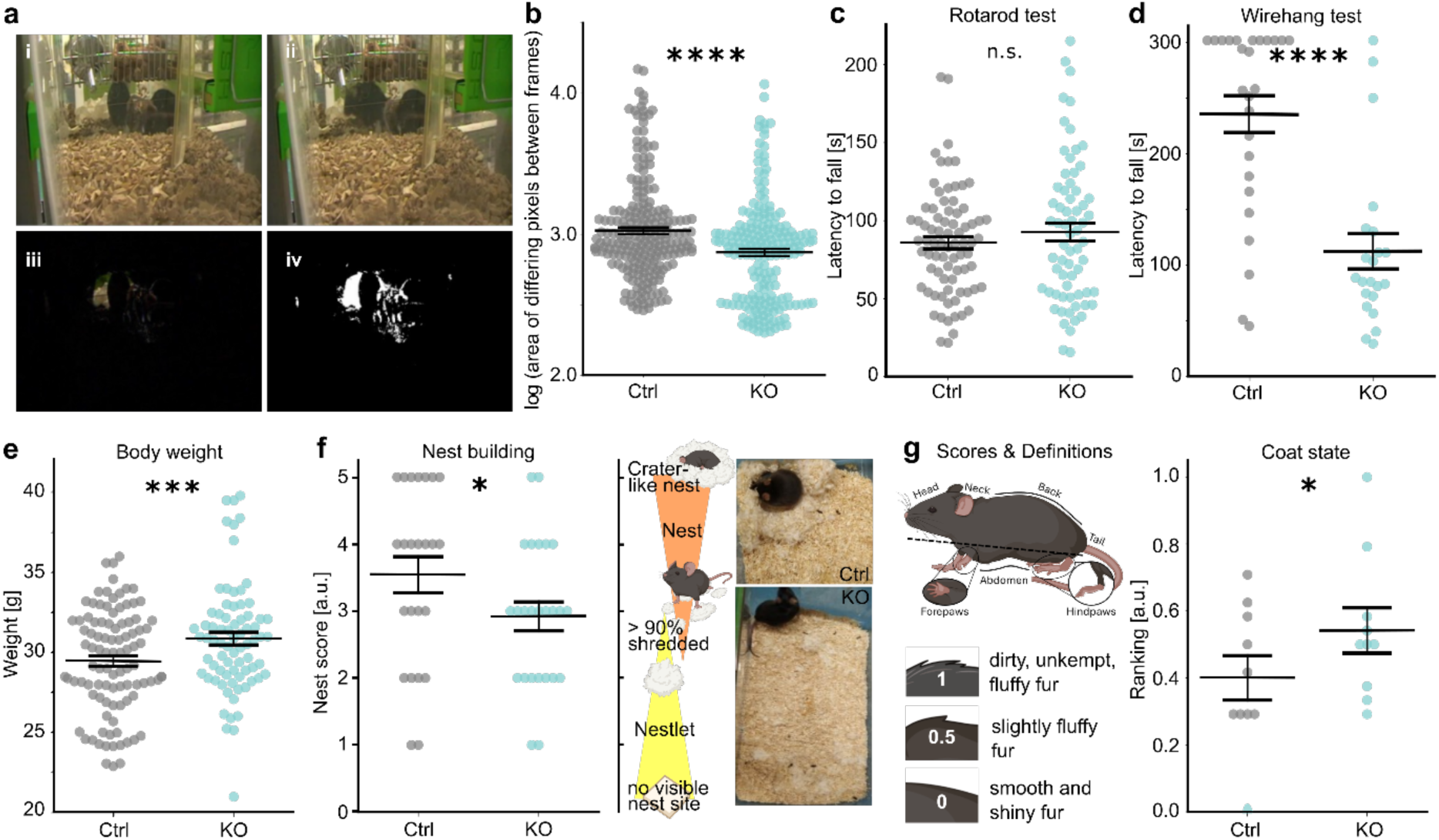
*Pvalb-Cre/tdTomato/Dnmt1 loxP^2^* (KO) mice display apathetic behavior. **(a–b)** Spontaneous activity within the home-cage was consistently diminished in KO mice compared to *Pvalb-Cre/tdTomato* (control/Ctrl) mice (*n* = 6 mice per genotype). **(a)** Mice were monitored in their home-cages to evaluate their spontaneous, undisturbed behavior. By comparing every fifth frame from the resulting videos (i, ii), the change in pixel values between frames (iii) was binarized (iv) to calculate the change in position of the contained animals. **(b)** Comparison of multiple 30-min time intervals revealed consistently lower activity levels of KO mice (Two-way ANOVA: *p_(Genotype)_* = 1.9×10^-5^; *p_(Time bin)_* = 0.98, *p_(Genotype x Time bin)_* = 0.91). **(c)** Average latency to fall from a rotating rod (‘Rotarod’) with increasing velocity showed no apparent motor deficits in knockout mice (*n*_(Control)_ = 13, *n*_(Knockout)_ = 11; ANOVA: *p* = 0.31). (**d**) A wire hang test showed a significantly reduced latency to fall in KO mice, which can indicate a difference in motor strength but is highly dependent on the animals’ body weight (*n*_(Control)_ = 13, *n*_(Knockout)_ = 11; ANOVA: *p* = 3×10^-6^). **(e)** Body weights were elevated for KO mice compared to controls (*n*_(Control)_ = 43, *n*_(Knockout)_ = 33; ANOVA: *p* = 0.0068). **(f)** Nest-building capacities were significantly decreased in KO (*n* = 25) versus control mice (*n* = 24; Generalized estimating equation model: *p_(Genotype) =_* 0.006). Nest quality was assessed using an ordinal scale from 1–5 as visualized by the schematic in the middle. As the representative images (right side) show, knockout mice were shredding the provided material but failed to assemble visible nest sites. **(g)** Assessment of the animals’ coat state as indicated in the illustration on the bottom right showed significantly higher rankings of the knockout mice, indicating worsened fur care (*n* = 12 mice per genotype; Mann-Whitney U test: *p* = 0.026). Panels b–g display individual data points with median (centre line) and interquartile range (upper and lower lines). Ctrl = control, KO = knockout; * *p* < 0.5, ** *p* < 0.01, *** *p* < 0.001, **** *p* < 0.0001.

Additional behavioral assays further supported an apathetic and anhedonic phenotype of *Dnmt1* KO mice. Nest-building, an innate behavior essential for heat conservation and shelter, is a sensitive readout of apathetic states in rodents^32^. Using a 1–5 ordinal scale, we found significantly lower nest scores in *Dnmt1* KO mice (Fig. 2f), typically shredding but failing to assemble the nesting material within 16 hours. Assessment of coat state, a marker of self-care and motivation^33,34^, also revealed systematic neglect in *Dnmt1* KO mice, with significantly higher cumulative scores compared to controls indicating a deterioration of their overall grooming and coat care (Fig. 2g).

Consistent with lower self-motivated activity, voluntary wheel running was strongly reduced (Fig. 3a), particularly during the dark phase, suggesting diminished motivation for rewarding behaviors^35,36^. KO males also displayed reduced sexual motivation, mounting females less frequently (Fig. 3b), and showing no attraction to female urine (Fig. 3c), although general social interaction remained intact (Fig. 3b). Taken together, *Dnmt1*-deficient mice show a cluster of behavioral traits, including apathy, anhedonia, and reduced spontaneous activity, that collectively reflect a dysphoric state resembling depressive symptoms.

**Figure 3:**
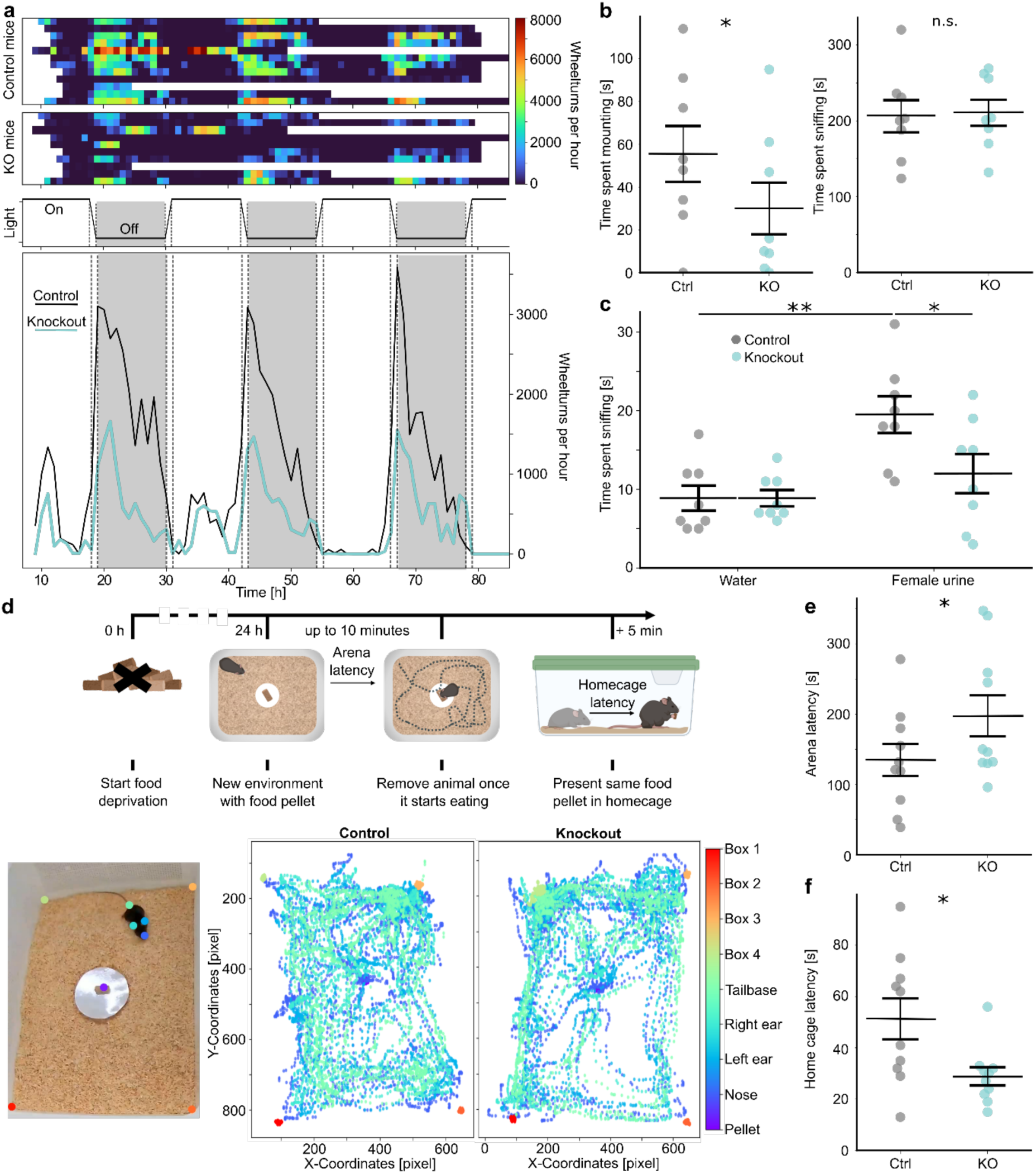
*Pvalb-Cre/tdTomato/Dnmt1 loxP^2^* (KO) mice display traits of an anhedonic, anxious phenotype. **(a)** Voluntary running on a wheel provided within the home-cage was strongly reduced in KO (*n* = 9) versus *Pvalb-Cre/tdTomato* (control/Ctrl) mice (*n* = 11). Running wheels were provided in the home cages of single-housed mice for up to three nights. Almost all mice displayed fluctuating utilization of the running wheel (upper heat maps) based on the daytime (middle graph), but in general, KO mice were running less frequently than controls. The overall reduced activity of knockout mice is discernible in the mean running time averaged per genotype (lower graph). **(b)** Evaluating the interaction of male KO and control mice (*n* = 8 mice per genotype) with female C57BL/6J-mice in a 10-min time frame revealed that KO males spent significantly less time trying to mount the females (left graph, Student’s *t*-test: *p* = 0.019), while they spent similar times sniffing their conspecifics (right graph, Student’s *t*-test: *p* = 0.44). **(c)** Presentation of pooled urine from female C57B/6J mice significantly increased time spent sniffing in control but not in KO mice (*n* = 8 mice per genotype; Linear mixed effects model: *p_(Genotype x stimulant type)_* = 0.048, Tukey post-hoc test: *p_(Ctrl: water vs urine)_* = 0.0017, *p_(KO: water vs. urine)_* = 0.25, *p_(Water: Ctrl vs KO)_* = 1.0, *p_(Urine: Ctrl vs KO)_* = 0.0413). **(d)** Conditional *Dnmt1* deletion in PV cells induces anxiety in a novelty-suppressed feeding (NSF) test (*n* = 10 per genotype). Upper row: Scheme and schedule of the NSF test. Mice were food deprived for 24 h before being transferred into an arena with new bedding and a centrally attached food pellet of known weight. Once the animals started chewing the presented food, their latency was recorded, and they were immediately removed to their home-cage. Subsequently, the pellet from the arena was presented in the home-cage and the animals’ latency to approach and consume it was recorded. Videos taken during the NSF test were labeled using DeepLabCut, to evaluate the animals’ behavior until they approached the food pellet (lower graphs). **(e)** KO mice displayed a significantly increased latency to approach the presented food pellet in the arena compared to controls, suggesting increased levels of anxiety. **(f)** In their home cages, KO mice approached the same food pellet significantly faster than control mice. Panels b, c, e, and f display individual data points with median (centre line) and interquartile range (upper and lower lines). Ctrl = control, KO = knockout; * *p* < 0.5, ** *p* < 0.01, *** *p* < 0.001, **** *p* < 0.0001.

### *Dnmt1* deletion in PV interneurons leads to increased anxiety in mice

To assess whether *Dnmt1* KO mice also display altered anxiety levels, we performed the novelty-suppressed feeding (NSF) test, which measures the conflict between food-seeking motivation and anxiety in a novel environment (Fig. 3d). After 24 hours of food deprivation, both genotypes showed similar weight loss (Extended Data Fig. 6a), implying a similarly strong motivation to obtain food.

Nevertheless, *Dnmt1* KO mice took significantly longer to approach and consume a food pellet centrally placed in an open arena (Fig. 3e) and foraged closer to the walls of the arena, reflected by an increased mean distance to the pellet (Fig. 3d). To confirm that this effect reflected anxiety rather than altered motivation, mice were transferred to their home-cages immediately after they started biting the pellet. With the food deprivation still serving as an active motivator in this familiar environment, KO mice approached the pellet faster than controls (Fig. 3f) and consumed equal amounts within 5 minutes (Extended Data Fig. 6b). These results demonstrate that DNMT1 loss in PV interneurons increases novelty-induced anxiety, further complementing the depression-like phenotype of KO mice.

### Single-nucleus transcriptomics reveals non–cell-autonomous glial remodeling

The combined electrophysiological and behavioral results show that DNMT1 loss in PV interneurons disrupts inhibitory control, weakens gamma synchrony, and gives rise to a spectrum of depression- and anxiety-related behaviors. Notably, DNMT1 loss not only increased firing of PV interneurons but also elevated spontaneous activity across the broader neuronal population, consistent with an adaptive network response to altered inhibition. These findings highlight the cell-autonomous role of DNMT1 in PV interneurons but also the need to understand how such local dysfunctions reverberate through cortical circuits.

Given that interactions between inhibitory and excitatory neurons are also tightly regulated by glia cells, we next turned to single-nucleus RNA sequencing to assess whether *Dnmt1* deletion in PV interneurons exerts non-cell-autonomous effects on all surrounding cell types. This approach enabled us to capture transcriptional adaptations across neurons and glial populations, providing a systems-level view of how DNMT1-dependent dysfunction in PV interneurons destabilizes cortical networks.

Unsupervised clustering of single nuclei isolated from the cortical tissue of adult *Dnmt1* KO and control mice revealed the expected diversity of neuronal and glial cell types, including excitatory neurons, PV and non-PV interneurons, oligodendrocytes, astrocytes, and microglia (Fig. 4a, b). Interestingly, although *Dnmt1* deletion was restricted to PV interneurons, relative proportions of excitatory subclasses were also altered. Population analysis revealed increased proportions of layer 2/3 intratelencephalic (IT) and layer 4 sensory neurons, combined with a marked reduction in layer 5/6 IT neurons (Extended Data Fig. 7). This underrepresentation of layer 5 neurons was also observed in earlier studies and likely reflects a combination of factors, including slower fixation and RNA degradation in deeper cortical layers, mechanical fragility of large nuclei with low nucleus-to-cytoplasm ratios, and inefficient recovery of long, chromatin-associated transcripts characteristic of layer 5 pyramidal cells^37–40^. Consequently, apparent shifts in excitatory subclass proportions in split-pool snRNA-seq datasets should be interpreted with caution, as they may partly arise from these technical biases rather than genuine biological differences^37^.

**Figure 4:**
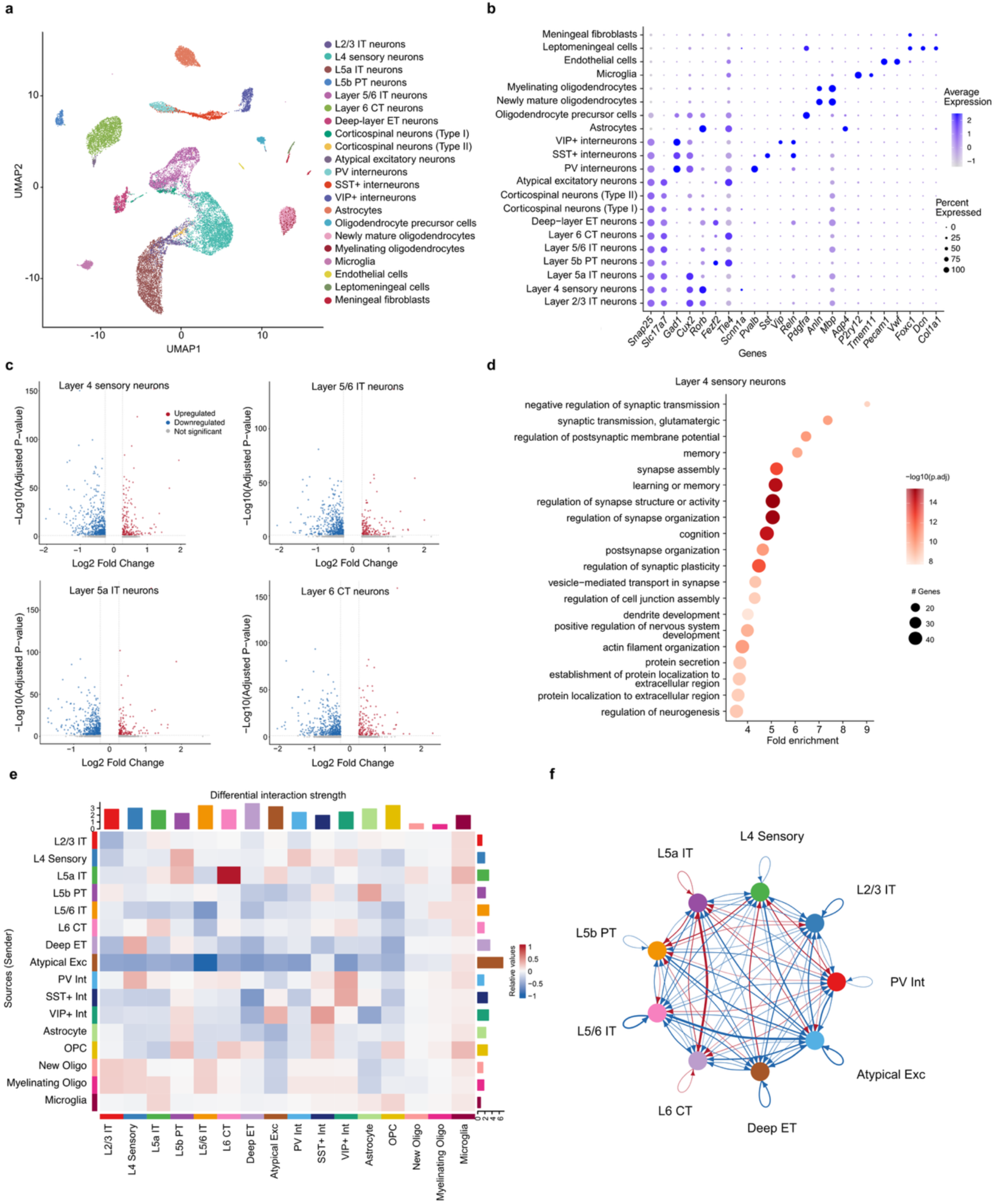
Single-nucleus RNA sequencing reveals transcriptomic shifts in excitatory neurons as well as a redistribution of interaction strength between PV interneurons and excitatory subtypes in *Pvalb-Cre/tdTomato/Dnmt1 loxP^2^* (KO) mice. **(a)** Uniform manifold approximation and projection (UMAP) embedding reveals the expected cellular composition of the neocortex. **(b)** Dot blot of marker genes across the identified clusters. **(c)** Volcano plots illustrate the extent of upregulated and downregulated genes in excitatory neurons from layers 4–6 of KO mice (log_2_FC ≥ 0.26, adjusted *p* value < 0.05). **(d)** Gene ontology analysis of differentially expressed genes (DEGs) in the layer 4 sensory neurons reveal an enrichment for biological processes related to synaptic transmission, organization, and activity. **(e)** CellChat-based mapping demonstrates altered interaction networks among neuronal and glial cell clusters in the KO cortex. The senders, i.e., the clusters the signaling event is initiated from, are listed in the y-axis while the clusters receiving the signal are found in the x-axis. **(f)** The circular network diagram depicts altered connectivity patterns between excitatory neurons and PV interneurons in KO mice. Increased and decreased communication probabilities in KO mice are depicted in red and blue, respectively.

We further detected broad and robust transcriptional alterations across multiple cell types between samples from control and KO mice, with prominent changes in excitatory neurons of layers 4–6 (Fig. 4c). Notably, the differentially expressed genes of excitatory neurons in layer 4, the cortex’s principal thalamorecipient layer which is also particularly populated by PV interneurons^41^, were enriched for gene ontology (GO) terms related to synaptic transmission, structure and organization. These results are consistent with altered input-layer gating due to disrupted inhibitory control after *Dnmt1* deletion (Fig. 4d). Since DNMT1 loss was selective to PV interneurons, these alterations likely reflect secondary, non-cell-autonomous effects.

A plausible mechanism is that the increased GABAergic transmission we observed electrophysiologically (Fig. 1b) directly affects the surrounding network through altered synaptic signalling. To systematically explore such effects, we next applied cell–cell communication analysis (CellChat) to identify changes in signaling pathways linking PV interneurons to surrounding neurons and glial populations. Differential interaction analyses showed redistributed communication strength and altered signaling pathways between PV interneurons and excitatory subclasses as well as glial populations in the *Dnmt1* KO mice (Fig. 4e, f).

Communication between excitatory neurons and PV interneurons showed a layer-specific redistribution. Outputs from PV cells were prominently increased toward layer 4 sensory neurons, aligning the strongest inhibitory reweighting with the primary sensory input gateway. Modest increases occurred toward layer 5/6 IT and layer 6 corticothalamic neurons, whereas projections to layer 2/3 IT and deep-layer extra-telencephalic neurons were slightly reduced (Fig. 4e, f). This laminar imbalance suggests a shift toward input-layer-centric inhibition that may favor rapid sensory gating but weaken distributed recurrent stability. Conversely, inputs from multiple excitatory neurons to PV interneurons declined, potentially explaining the reduced inhibitory efficacy despite PV hyperactivity (Fig. 4e, f).

The decline in inputs to PV interneurons suggests a potential compromise in the integrity of perineuronal nets (PNNs), a specialized extracellular matrix structure that is critical for stabilizing synaptic contacts onto PV cells and supports their high-frequency firing^14,15^. As PNNs require continuous support from the oligodendrocyte lineage, we next looked at their interactions with PV interneurons. Although overall oligodendrocyte lineage proportions were unchanged (Fig. 5a), we detected a decoupling of the PV–oligodendrocyte precursor cells (OPC) axis with decreased signaling in both directions as well as a selective reduction in newly mature *Pdgfc*-positive perineuronal oligodendrocytes^42,43^ that synthesize, organize, and anchor extracellular matrix (ECM) constituents (Fig. 4d, Fig. 5b). Concordantly, perineuronal oligodendrocytes in *Dnmt1* KO mice downregulated genes implicated in stress-resilience, ECM-remodeling, and structural programs (*Sgk1*, *Fkbp5*, *Bcl2l1*, *Htra1*, *Mobp*, *Zbtb16*), indicating a reduced capacity to sustain high-demand perisomatic coverage and ECM organization (Extended Data Fig. 8). Notably, PV interneuron–oligodendrocyte and reciprocal oligodendrocyte–PV connections were significantly reweighted in *Dnmt1* KO mice, indicating a disruption in PV–oligodendrocyte coupling (Fig. 5b–d).

**Figure 5:**
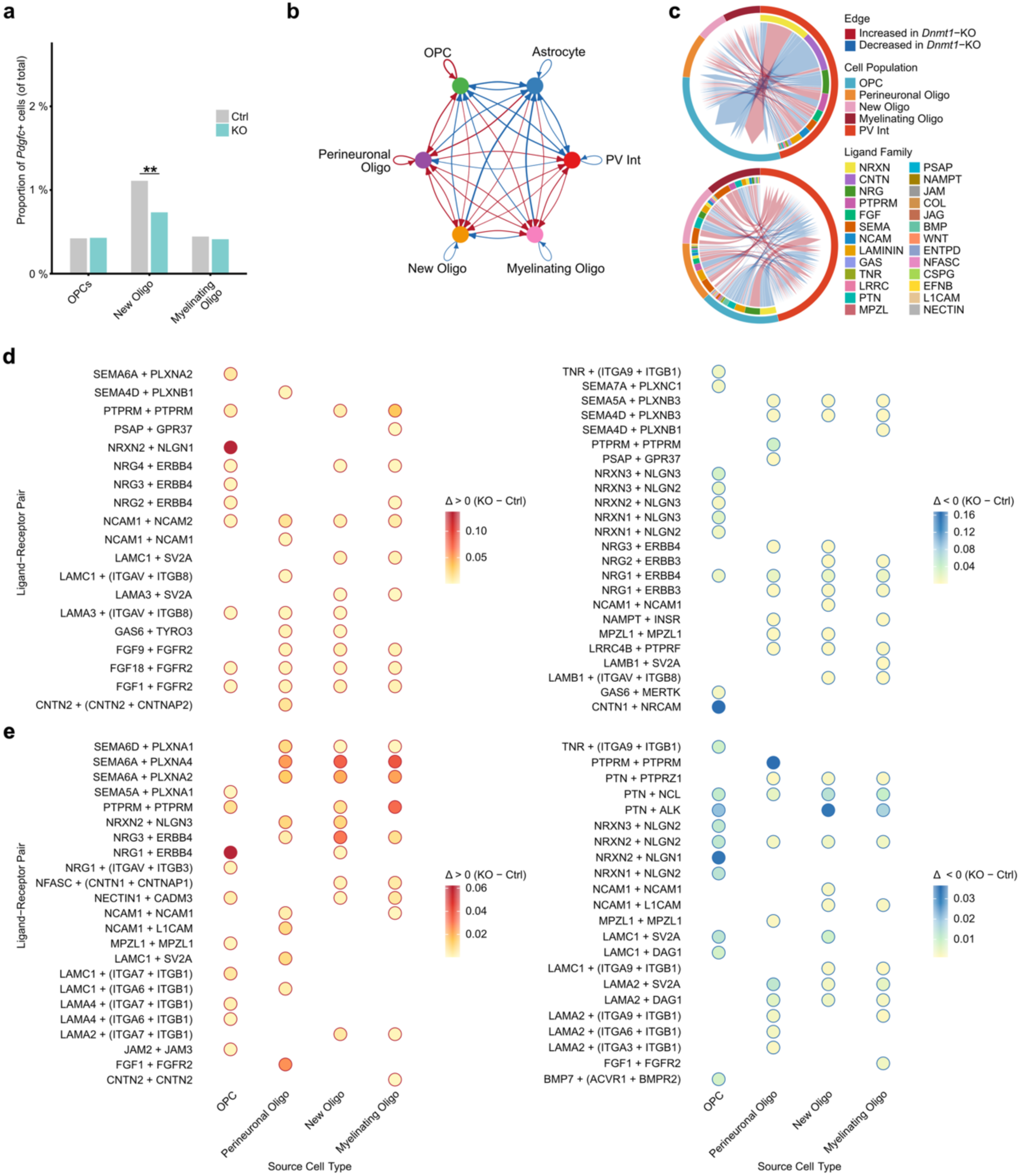
DNMT1 deficiency in PV interneurons reshapes PV-oligodendroglial communication and alters ligand–receptor interactions crucial for perineuronal nets (PNNs) in *Pvalb-Cre/tdTomato/Dnmt1 loxP^2^* (KO) mice. **(a)** The *Pdgfc*-expressing population within newly mature oligodendrocytes, representing perineuronal oligodendrocytes, is significantly reduced in the KO cortex (two-sample proportion test, *p* = 0.005). **(b)** The circular network diagram illustrates differential interactions between oligodendroglial cells and PV interneurons in KO mice. Increased and decreased communication probabilities in KO mice are depicted in red and blue, respectively. **(c)** Chord diagrams depict the altered interactions grouped by ligand/receptor family, separated based on directionality. Upper diagram: PV interneurons → oligodendroglial cells; lower diagram: Oligodendroglial cells → PV interneurons. Ligand–receptor pairs with known roles in stabilizing PNNs, the perisomatic niche and cell–cell interactions have altered communication probability in **(d)** outgoing signaling from PV interneurons to oligodendroglial cells as well as **(e)** incoming signaling from oligodendroglial cells to PV interneurons.

Therefore, we next examined which specific ligand–receptor pairs contributed most strongly to these altered communication probabilities. Outgoing signaling from PV interneurons to oligodendroglial cells and incoming signaling from oligodendroglial populations to PV interneurons are shown in Figure 5d and 5e, respectively. Overall, more extensive alterations were detected in the incoming direction, suggesting that PV interneurons become particularly sensitive to glia-derived cues when DNMT1 is lost. In contrast, PV-to-glia signaling showed fewer but distinct changes, mainly involving PV-OPC interactions. Specifically, NRXN2–NLGN1 signaling was increased in *Dnmt1* KO samples, whereas CNTN1–NRCAM communication was reduced, implying weakened contact-mediated coupling^44^.

Among glia-to-PV pathways, several axes were markedly affected across OPCs and mature oligodendrocyte subtypes. NRG1–ERBB4 signaling from OPCs to PV interneurons was significantly diminished, indicating a loss of trophic support normally promoting PV excitability and metabolic stability^45,46^. In contrast, SEMA–PLXN signaling was enhanced, particularly from differentiated oligodendrocyte populations, potentially reflecting a structural remodeling response. Notably, semaphorins have been reported as constituents of PNNs^47^. PTPRM–PTPRM signaling from perineuronal oligodendrocytes to PV cells, as well as NRXN–NGLN signaling from OPCs, was reduced (Fig. 5e). PTPRM can mediate homophilic interactions, known to be involved in cell-cell adhesion and synapse formation^48^ while NRXN–NGLN signaling operates as an axo-glial communication pathway between axonal neurexin and oligodendrocytic neuroligin. NGLN3 is expressed in oligodendrocyte precursor and mature cells, where it interacts with axonal NRXN to promote myelin formation, maintenance, and oligodendrocyte differentiation^49^. Together with the reduced CNTN1–NRCAM interactions (Fig 5d) and attenuated TNR–integrin inputs onto PV somata (Fig. 5e), these changes indicate deficiencies in perisomatic ECM anchoring and PNN cohesion^50–52^. Collectively, these alterations indicate a reorganization of neuron-glia communication that compromise PNN stabilization and PV synaptic integration, leading to a reduction in inhibitory efficacy of PV interneurons.

### Histological validation confirms perineuronal net deficits

To validate the results of the cell-cell interaction analyses, we next aimed to visualize potential changes in the PNN formation. Indeed, consistent with the predicted disruptions in neuron–glia signaling, histological analysis revealed a reduction of Wisteria floribunda agglutinin (WFA)-labeled PNNs in the *Dnmt1* KO mice (Fig. 6a–d, Supplementary Video 1, 2). This points to a disruption of the structural support mechanisms that normally stabilize PV interneuron function. While the fraction of PV cells enwrapped by PNNs remained unchanged, WFA labeling intensity was significantly decreased (Fig. 6e, f). Given that PNNs stabilize PV firing properties and gamma synchrony^15,53^, these findings implicate glial and extracellular matrix abnormalities as amplifiers of PV interneuron deficits, contributing to both the physiological (reduced PV efficacy, weakened gamma) and behavioral (apathy/anhedonia/anxiety) phenotypes in *Dnmt1* KO mice.

**Figure 6:**
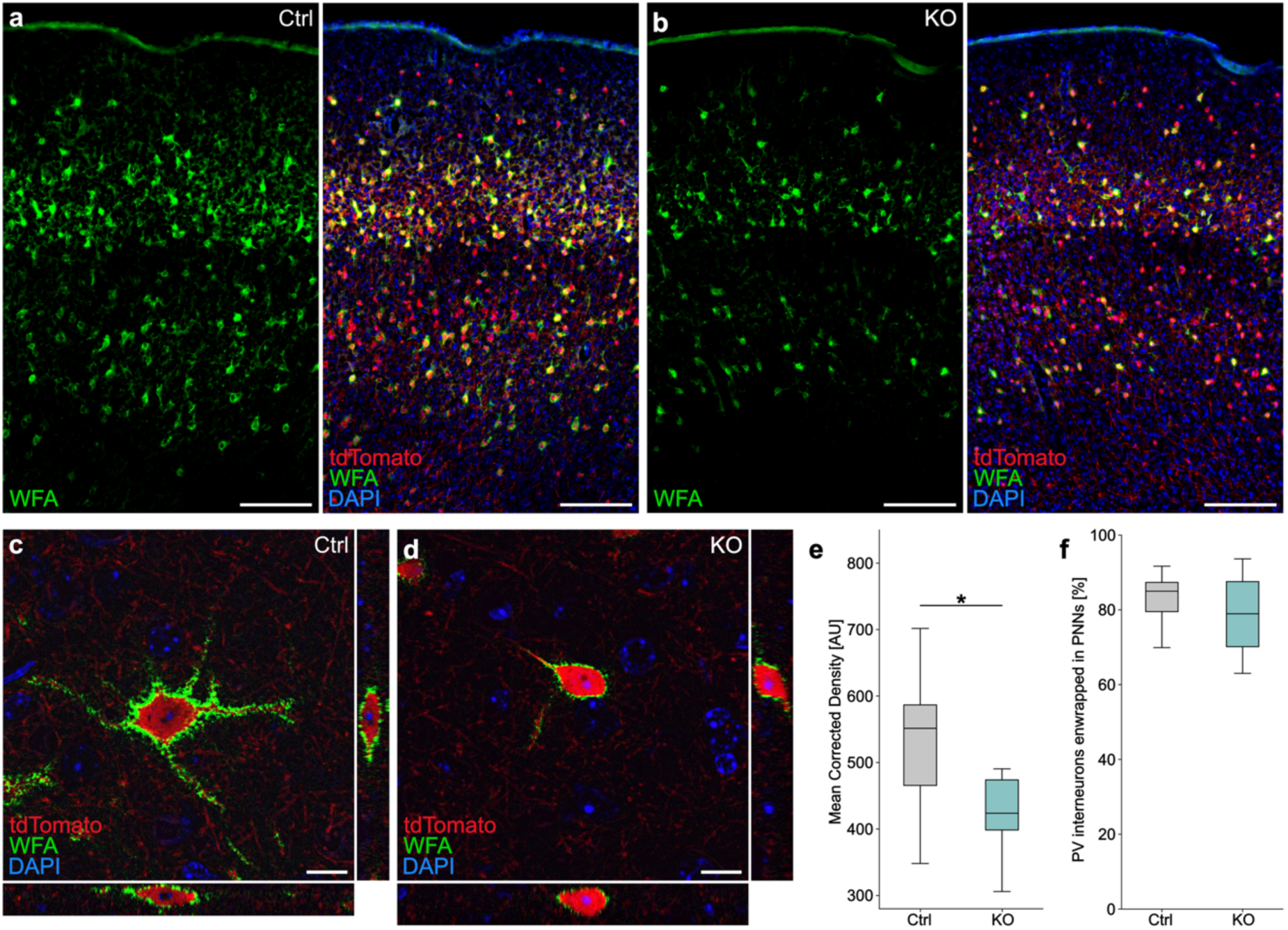
Loss of DNMT1 in PV interneurons disrupts perineuronal nets (PNNs) in the neocortex of *Pvalb-Cre/tdTomato/Dnmt1 loxP^2^* (KO) mice. **(a–b)** Representative microphotographs of the primary somatosensory cortex (S1) in coronal brain sections of *Pvalb-Cre/tdTomato* (Ctrl; **a**) and *Pvalb-Cre/tdTomato/Dnmt1 loxP^2^*(KO, **b**) mice stained for PNNs using Wisteria floribunda lectin (WFA). Scale bars: 200 µm. **(c–d)** Orthogonal XZ und YZ projections show the three-dimensional localization of a PV interneuron soma and surrounding PNNs stained with WFA. Scale bars: 10 µm. **(e)** Mean corrected WFA signal density is reduced in the neocortex of KO mice (box plots: median, interquartile range, min-to-max; paired Student’s *t*-test, *p* = 0.046; *N* = 3 biological replicates with *n* = 3 slices each per group). **(f)** Proportion of PV interneurons enwrapped by PNNs in S1 of KO mice is unchanged (box plots: median, interquartile range, min-to-max; paired Student’s *t*-test, *p* = 0.277; *N* = 3 biological replicates with n = 8 slices per genotype). * *p* < 0.05.

Together, these findings provide a mechanistic bridge from DNMT1 loss in PV interneurons to disrupted inhibitory signaling, defective PV-glia communication and impaired PNN integrity, ultimately destabilizing the cellular ecosystem required for cortical network balance and gamma synchrony.

## Discussion

Here we show that deleting *Dnmt1* specifically in cortical PV interneurons reshapes inhibition across multiple scales, including cellular, network, and behavior, culminating in depression- and anxiety-like phenotypes. In vivo Neuropixels recordings revealed increased spontaneous firing of PV cells and elevated baseline activity in surrounding cortical populations, accompanied by a reduced inhibitory impact of PV activation on local circuits. The dominant consequence of *Dnmt1* deletion is therefore not reduced PV interneuron activity but diminished synaptic efficacy and a corresponding loss of inhibitory control. This is consistent with our previous finding that *Dnmt1* loss enhances GABAergic vesicle replenishment and release at PV terminals^13^. We interpret the global rise in excitatory neuron firing and the blunted response to PV photostimulation as a homeostatic network adaptation to counteract persistently elevated GABAergic drive by down-weighting inhibitory influence, a phenomenon observed in vitro and in vivo when networks strive to stabilize E/I balance^54^. Moreover, because PV interneurons form GABAergic autapses that regulate their fast-spiking dynamics and contribute to gamma synchrony^55^, the observed increase in PV firing may also represent an adaptive response to altered self-inhibition. These network adjustments likely contribute to the smaller, delayed, and less precise visually evoked responses and the loss of a feedback-inhibitory rebound across layers.

Despite these physiological changes, basic perceptual and learning capacities remained intact, suggesting a circuit-level compensation potentially via subcortical pathways that sustain task performance even when cortical precision is reduced. Nevertheless, DNMT1 loss degraded PV-dependent network dynamics across sensory (V1, S1) and prefrontal (ALM) cortical areas, PV-mediated inhibition was weakened, and visually evoked gamma oscillations were markedly reduced, in line with the central role of PV cells in generating gamma rhythms^21,22^.

*Dnmt1* KO mice exhibited an apathy- and anhedonia-like phenotype, including reduced home-cage activity, poor nest construction, deteriorated coat condition, diminished wheel running and sexual motivation, and prolonged latency in novelty-suppressed feeding. Food intake and circulating ghrelin, insulin, and PYY levels remained unaltered (Extended Data Fig. 4a–d), whereas body weight and leptin were elevated (Fig. 2e, Extended Data Fig. 4d), indicating reduced energy expenditure rather than primary hypothalamic dysregulation. Correspondingly, PV cells in the arcuate nucleus showed no overlap with POMC and only partial overlap with AgRP populations. These findings point to cortical, rather than hypothalamic, mechanisms underlying the dysphoric phenotype, in line with evidence linking PV interneuron dysfunction and impaired gamma synchrony to mood disorders in both humans and rodent models^11,22,27,56,57^.

A central point of this study is that a PV-restricted, cell-autonomous manipulation of *Dnmt1* expression propagates across the network, leading to widespread, non-cell-autonomous alterations. Single-nucleus RNA-seq uncovered a remodeling of excitatory neuron composition as well as their transcriptional rewiring of genes involved in synaptic structure and transmission. Consistent with these molecular changes, circuit-level analyses demonstrated a laminar redistribution of inhibitory signaling. Communication between excitatory neurons and PV interneurons showed a marked layer-specific reorganization, with PV outputs prominently reweighted toward layer 4 while connections to layer 2/3 and deep-layer extratelencephalic neurons were modestly reduced. Such preferential strengthening of feedforward inhibition within the sensory input layer resembles canonical thalamocortical PV architectures, where fast-spiking PV cells mediate rapid inhibitory gating of sensory signals^58,59^. Modest increases in PV targeting of layer 5/6 subtypes may reinforce corticothalamic feedback loops, whereas diminished supragranular inhibition could weaken recurrent excitatory–inhibitory stability across cortical columns^60,61^. Interestingly, excitatory drive onto PV cells was reduced, in line with the notion that PV interneurons can become hyperactive yet functionally decoupled when excitatory afferents weaken, resulting in ineffective inhibition despite elevated firing^56,62^. Collectively, these laminar rebalancing effects suggest an input layer-centric inhibitory bias that overemphasizes sensory gating at the cost of distributed network stability.

Similarly, communication between PV interneurons and oligodendroglial cells was particularly reweighted in the cortex of KO mice, accompanied by a selective depletion of *Pdgfc*-expressing perineuronal oligodendrocytes and reduced expression of genes maintaining ECM integrity. Oligodendrocyte lineage cells are crucial signaling partners for interneurons, responding dynamically to inhibitory signals via GABA receptors^63^. The coordinated loss of key ligand–receptor interactions between PV cells and oligodendroglial cells, such as TNR–integrin, and NRXN–NLGN, points to a breakdown of perisomatic adhesion and PNN cohesion, leading to impaired network connectivity and plasticity^14^.

Genetic deletion of TNR disrupts PNN morphology, causing irregular net structuring around cortical interneurons and altered conduction velocity, highlighting its importance for cohesive perisomatic ECM^64^. Integrin receptors, in turn, contribute to synaptic stabilization by binding ECM components such as TNR, reinforcing PNN architecture and modulating excitatory–inhibitory balance^17^. On the synaptic adhesion side, neurexins (NRXNs) and neuroligins (NLGNs) form transsynaptic complexes that coordinate synapse formation and maintain structural adhesion and communication between neurons^65,66^. Perturbations in NRXN–NLGN signaling have been shown to destabilize synaptic alignment and weaken the structural cohesion of perisomatic synapses^67,68^. Moreover NRXN–NGLN signaling mediates axo-glial communication by coupling presynaptic neurexins on axons to neuroligin-3 (NGLN3) on oligodendrocyte lineage cells. NGLN3 is expressed in both precursor and mature oligodendrocytes, where its interaction with axonal NRXNs facilitates differentiation, myelin formation and stability^49^.

As a consequence of these disrupted neuron-glia interactions, elevated PV activity and altered spiking patterns may drive activity-dependent glial plasticity that reshapes PNN and neuro-glial positioning. Histological validation confirmed this systems-level implication, showing diminished net intensity of WFA-labeled PNNs in KO mice. This is consistent with the established role of PNNs in maintaining PV interneuron excitability, synaptic integration, and gamma synchrony^14^. Indeed, PNN degradation has been shown to weaken gamma oscillations and cognitive precision^14,69^, providing a mechanistic link between synaptic inhibition and network disruptions.

Because PNNs stabilize PV excitability and support oscillatory coherence, their loss provides a functional bridge between reduced PV efficacy, weakened gamma power, and mood-related behavior. We propose a feed-forward model in which DNMT1 loss increases presynaptic GABA release, triggering homeostatic desensitization of inhibitory signaling and glial remodeling that culminates in ECM degradation and network desynchronization. The resulting disruption of PNNs may further destabilizes PV function, amplifying network desynchronization and dysphoric behavior.

This cascade reconciles diverse GABAergic findings in mood disorders: alterations in PV efficacy, gamma coherence, and PNN integrity can occur without cell loss, reflecting broader neuron-glia-ECM dysregulation. By positioning DNMT1 as an upstream regulator of this axis, our results highlight an epigenetic mechanism by which inhibitory microcircuits coordinate with myelination and extracellular scaffolds to sustain cortical timing.

## Methods

### Animals

Transgenic *Pvalb-Cre/tdTomato/Dnmt1 loxP^2^*-mice were used as knockout animals and *Pvalb-Cre/tdTomato*-mice as controls. Unless stated differently, all experiments were conducted using male mice aged 3–6 months. Both mouse lines were established on a C57BL/6-background by crossing in the *Pvalb-Cre* line (obtained from Christian Huebner, University Hospital Jena, Germany and described in Hippenmeyer et al.^70^) and the *tdTomato* transgenic reporter mouse line (obtained from Christian Hübner, University Hospital Jena, Germany and described in Madisen et al.^71^). To obtain *Dnmt1*-knockout mice, *Dnmt1* floxP^2^ mice (B6;129Sv-Dnmt1^tm4Jae^/J, Jaenisch laboratory, Whitehead Institute; USA; Jackson-Grusby et al.^72^) were crossed in, in which LoxP-sites are flanking exons 4 and 5 of the *Dnmt1* gene. This results in a null *Dnmt1* allele upon Cre-mediated excision^72^. Correct genotypes were validated with the following primers: *Dnmt1* forward 5′-GGGCCAGTTGTGTGACTTGG and reverse 3′-CCTGGGCCTGGATCTTGGGGA (results in a 334 bp wildtype and 368 bp mutant band); *tdTomato* wildtype forward 5′-AAGGGAGCTGCAGTGGAGTA, wildtype reverse 3′-CCGAAAATCTGTGGGAAGTC, mutant forward 5′-CTGTTCCTGTACGGCATGG and mutant reverse 3′-CTGTTCCTGTACGGCATGG (results in a 297 bp wildtype and 196 bp mutant band); *Pvalb*-*Cre* forward 5′-AAACGTTGATGCCGGTGAACGTGC and reverse 3′-TAACATTCTCCCACCGTCAGTACG (resulting in a 214 bp fragment). All animal procedures were performed in strict compliance with the EU directives 86/609/EWG and 2007/526/EG guidelines for animal experiments and were approved by the local governments (Thueringer Landesamt, Bad Langensalza, Germany and Landesamt für Natur, Umwelt und Verbraucherschutz Nordrhein-Westfalen, Recklinghausen, Germany). Unless indicated otherwise, animals were housed under standard housing conditions: mice were kept in compatible groups, under 12 h light/dark conditions, with ad libitum access to food and water, and nestlets and tunnels as environmental enrichments. The weight of individual animals was determined prior to behavioral experiments or prior to sacrificing and brain preparation.

### Electrophysiological recordings

#### Surgeries

For all surgeries isoflurane in oxygen was used as anesthesia (3% isoflurane for induction, 1–2% for maintenance), and subcutaneous injections of carprofen (4 mg/kg, Rimadyl, Zoetis GmbH) and buprenorphine (0.1 mg/kg, Buprenovet sine, Bayer Vital GMBH) for analgesia. Eye ointment (Bepanthen, Bayer Vital GmbH) was applied to their eyes to prevent damage from drying out. After medially incising the dorsal skin of the head, it was pushed outward and fixed in place utilizing tissue adhesive (Vetbond, 3M). Dental cement (C&B Metabond, Parkell; Ortho-Jet, Lang Dental) was applied on top of the tissue adhesive to create a support structure for a custom-built circular headbar.

Viral injections (AAV1.shortCAG.dlox.hChR2(H134R).WPRE.hGHp, Zurich Vector Core, viral titer = ∼8×1012 vg/ml) were carried out according to the experimental scheme. In naive mice (Fig. 1a–I, 2a–c), S1 and V1 were targeted unilaterally, whereas in trained animals (Fig. 2g–i, Extended Data Fig. 2b, c) bilateral injections in ALM were conducted. The positions of all injections were chosen by stereotactic coordinates (ALM: AP 2.5/ ML −1.5; V1: AP −4/ ML −2; S1: AP −2/ ML −3.5). Injections were conducted using a thin glass pipette with a micropump (Nanoject3, Drummond Scientific) after thinning out the skull with a dental drill. At each site, two quantities of 200 nl were injected at different cortical depths (300/600 µm), adding up to a total volume of 400 nl virus per target region. The 200 nl were injected in 20 steps using 10 nl portions at 14 s intervals. For electrophysiological recordings, circular craniotomies were performed using a biopsy puncher and then covered with a round coverslip of similar size using light-curable dental cement (DE Flowable composite, Nordenta) to fix it in place. The coverslips contained a small opening to later allow access for the Neuropixels probe. Between recordings, this opening was covered with silicon. A thin layer of cyanoacrylate (Zap-A-Gap CA+, Pacer technology) was used to cover the skull. After the surgery, buprenorphine (0.1 mg/kg body weight) was again injected subcutaneously. The animal’s home cage was kept on a heating pad following the surgery until the mouse recovered from anesthesia. To prevent inflammation of the surgical wounds, the animals received a subcutaneous injection of carprofen on post-op days 1 and 2 while the animal’s drinking water was supplemented with antibiotics (enrofloxacin 0.025 mg/ml) and analgesics (buprenorphine 0.0125 mg/ml) for at least three days. Progression of wound healing and the animal’s general health were scored daily following the surgery.

To record from naive animals, two mice per genotype were utilized; all of them were approx. 6 months of age during surgery. For behavioral analysis and recordings in trained mice, 7 animals (3 control/4 knockout) were used; they were approx. 8 weeks old during surgeries.

### Neuropixels recordings

Electrophysiological recordings were carried out in head-fixed mice using Neuropixels 1.0 probes. Recordings were carried out on four consecutive days. Probes were painted with DiD cell labeling solution (Invitrogen V22887) before each recording to later identify recording sites in fixed brain slices.

The stimulation protocol was started 5–10 min after insertion and we recorded high pass-filtered data above 300 Hz at 30 kHz and low pass filtered signals between 0.5–100 Hz at 2.5 kHz from the bottom 384 channels of the Neuropixels probe (∼3.8 mm active recording area). Signals were acquired with an external Neuropixels PXIe card (IMEC, Belgium) used in a PXIe-Chassis (PXIe-1071, National Instruments, USA). Triggers and control signals for different stimuli were separately recorded as analogue and digital signals using the SpikeGLX software (Janelia Farm Research Campus, USA; Bill Karsh, https://github.com/billkarsh/SpikeGLX).

The visual stimulus to test sensory responses in V1 and ALM consisted of a 0.1-s long full-field low-pass filtered Gaussian noise patterns with a cutoff at 0.12 cycles per degree and a temporal frequency of 1 Hz^73^. In each trial, the stimulus was presented twice, with an inter-stimulus interval of 0.5 s. Visual stimulation to induce gamma oscillations consisted of 5-s-long full field drifting square wave gratings at horizontal orientation and a spatial frequency of 0.04 cycles per degree. All visual stimuli were presented at 17-cm distance from the right eye on a gamma-corrected LED backlit LCD monitor (Viewsonic VX3276-2K-MHD-2, 32”, 60Hz refresh rate). The overall stimulation protocol consisted of 50 trials of each type, presented in randomized order with inter-trial intervals between 3–5 seconds.

For optogenetic stimulation, we used a fiber-coupled laser system (ReadyBeam Bio1, FISBA AG), connected to a 105 µm glass fiber (0.22 NA) that was positioned over the cortical recording site. Optogenetic stimulation consisted of a 3-s long pulse at 488 nm with a peak power of 10 mW from the fiber tip. To avoid transient changes in PV activity, the pulse intensity was first increasing for 1 second, then remained at a peak intensity for 1 second, and lastly decreased back to zero for 1 second. The sequence of visual, optogenetic, and combined trials was fully randomized. For optotagging of PV interneurons, we presented 20 optogenetic stimulus sequences at 5 Hz for 1 second, with an individual pulse length of 10 ms and a peak power of 2 mW from the fiber tip. PV interneurons were then identified by short-latency responses (<2 ms) to individual laser pulses and an increase in firing rate during the 3-s-long optogenetic stimulation period.

To extract spiking activity, channels that were broken or outside of the brain were detected and removed from the recording using the Spike Interface analysis toolbox^74^. The remaining channels were then time-shifted and median-subtracted across channels and time. Corrected channels were then spike-sorted with the Kilosort 2.5 software and manually curated using phy (https://github.com/cortex-lab/phy). All sorted spiking and local field potential data were then analyzed using custom Matlab code (2020b, MathWorks).

Peri-event time histograms (PETHs) for each cluster were computed with a bin size of 10 ms and baseline-subtracted by the mean activity within 1 second before the laser onset. Trial-averaged local field potentials (LFPs) in response to visual stimulation were similarly baseline-corrected. To compute current source densities (CSDs) from LFP signals, we used the inverse CSD method by Pettersen et al.^75^. We applied the spline iCSD method, assuming a smoothly varying CSD between electrode contacts, based on polynomial interpolation. We assumed a homogeneous, isotropic conductivity of σ = 0.3 S/m within and directly above cortex^75^. To reduce spatial noise, the estimated CSD was subsequently convolved with a Gaussian spatial filter with a standard deviation of 0.1 mm. For LFP and CSD analyses, the signals from neighboring contacts at the same depth were averaged together to improve signal quality.

### Behavioral Assays

#### Evidence accumulation task

After full recovery from surgeries, animals were water-deprived for one day before introducing them to the behavioral task. To obtain rewards, mice needed to accumulate visual evidence in form of visual grating-like cues and indicate the side with the higher number of cues. Experiments were conducted in a custom-built setup with stimulus presentation, lick-detection and water reward controlled by a microcontroller (Teensy 3.2, PJRC) and Python code (Python 3.7). In general, the task consisted of three periods: 3 s sensory stimulation with randomized sequences of up to 6 cues on one or both sides, 0.5 s delay and a 2-s response window. In between trials was a 3.5-s inter trial interval, which was prolonged after missed trials or incorrect responses. Mice first learned to reliably detect visual stimuli, by indicating the side were all 6 cues (spatial frequency of 0.018 cpd, temporal frequency of 2 Hz) were shown. After mice performed more than 75% of trials correctly in 3 out of the last 4 sessions, distractor stimuli were added on the non-rewarded side. The number of cues on both the target and distractor side varied, resulting in different levels of difficulty depending on the absolute difference in cues shown on either side. Training was conducted using 3 control and 4 knockout animals.

### Morris water maze (MWM)

Visual navigation and learning capacities were investigated using an MWM. The setup consisted of a cylindric water-filled pool (Ø 1 m) surrounded by an opaque curtain, four distinct landmarks on the pool wall, and a transparent platform (Ø 10 cm). The setup was divided into four virtual quadrants with the platform being consistently in the same quadrant. The start position of the mouse was pseudo-randomly alternated between trials. On the first day of testing, mice started once per quadrant for habituation. The platform slightly protruded from water level to facilitate quicker locating. Upon finding the platform, the animal was required to remain on top of it for 5 s for the trial to be counted successful. If an animal did not find the platform or left it after less than 5 s and did not return onto it within the 90-s period of the trial, the mouse was manually guided to the platform before taking it back to their cages. Between trials, mice were kept under red light for at least 10 min. For the actual learning task, the platform was submerged (1–2 cm) and the maximum duration per trial was set to 60 s. Each mouse started three times per quadrant in a pseudo-random order, equaling 12 trials per day. This training scheme was conducted on five consecutive days. Mice were recorded from above and behavioral data were processed using ANY-maze (Stoelting Co., Wood Dale, USA). We tested 8 control and 8 knockout mice.

### Home cage activity

To assess the animals’ activity in their home cages, cameras were set up in front of cages of group-housed animals. Each cage was filmed on several separate days with a framerate of 25 frames per second. From each of these recordings, eight uninterrupted 30 min periods were used for further analysis. For each time period, a pixel-wise comparison with ImageJ was utilized to calculate the difference between every fifth frame of a video. For this, we used the ImageJ Image Calculator and adjusted the threshold to exclude background noise. Subsequently, we measured the area of differing pixels. We filmed 3 cages per genotype housing 2 mice each (6 control/6 knockout).

### Rotarod

A rotarod was used to assess balance and motor performance of mice. Initially, the mice (13 control/11 knockout) were individually habituated for 4 min at a constant speed of 4 rpm. The subsequent test phase comprised three consecutive trials, each lasting a maximum of 5 min, with continuous acceleration from 4 to 40 rpm.

### Wire hang test

To record neuromuscular strength, mice were placed on a metal wire cage lid, which was then slowly inverted and positioned over a type IV Makrolon cage (Fig. 3d). Each mouse was given three trials of max. 5 min, with inter trial intervals of approx. 3–4 min. In total, 24 mice (13 control/11 knockout) conducted this experiment.

### Food intake

Food intake of group-housed mice was quantified by assessing the change in cumulative pellet weight after either 24 h or 72 h. Animals remained in their familiar cage settings and pellet weight measurements were calculated relative to the number of animals per cage as the consumed amount per day. Tests were conducted on 4 control and 4 knockout cages (each cage containing 2 mice: 8 control/8 knockout). For each cage two 24-h periods and eight 72-h periods were recorded.

To measure food and water intake in single-housed mice, mice were kept individually for at least two weeks, before measuring pellet and water bottle weight of each cage on five consecutive days. For this, we used 4 control and 6 knockout mice.

### Nest-building test

Nest-building tests were executed as described by Deacon^78^ and as visualized in Fig. 3f. In brief, mice were weighed and transferred to individual testing cages (IVC cages, Tecniplast) with ad libitum access to food and water, but without any additional environmental enrichment. One nestlet of known weight was placed in each cage. The test was initiated at 5 pm and nest-building skills were scored the next morning at 9 am. Nests were scored according to the following criteria: Score 1 = no noticeable interaction with the nestlet (more than 90% of the nestlet remained intact), Score 2 = nestlet was partially torn (50–90% of the nestlet remained intact), Score 3 = nestlet was largely torn, but no identifiable nest site was build (10–50% of the nestlet remained intact, but less than 90% of shredded material is gathered within a quarter of the cage area), Score 4 = nestlet was used to build a flat nest (less than 10% of the nestlet remained intact, less than 50% of the nest circumference is build higher than the resting mouse), Score 5 = nestlet was used to build a (near) perfect nest (less than 10% of the nestlet remained intact, more than 50% of the nest circumference is build higher than the resting mouse). Each mouse was tested twice with tests being one week apart from each other. In the time between tests, mice remained single-housed without access to further enrichments beside the handed nestlet. Overall, 49 mice (24 control/25 knockout) at ages between 3–6 months were tested.

### Coat state

The animals’ coat state was assessed in a blinded manner. The scoring system was adapted from Nollet et al.^34^ as visualized in Fig. 3g. Seven different regions of the animal’s body were scored individually with one of three distinct ranks. In case of a good coat state (smooth and shiny fur, no tousled or spiky patches) rank 0 was assigned. Strongly matted, oily or even stained fur was assigned with a rank 1. Slight deviations from the optimum coat state were ranked with a 0.5. Among the investigated areas were both dorsal (head, neck, back, tail) and ventral areas (fore paws, abdomen, hind paws). Overall, 12 control and 12 knockout mice (2–7 months) were investigated.

### Voluntary wheel running

Single-housed mice were kept in cages with access to running wheels (TSE Systems). Animals were housed for up to three days in these cages. While food and water were provided ad libitum, no further enrichments despite the running wheels were provided. Each animals’ wheel usage was continuously recorded and subsequently evaluated using a custom-written Python script. In total, 11 control and 9 knockout mice were utilized.

### Interaction test

To test for social interest in female conspecifics, virgin C57BL6J-females were transferred to cages of virgin transgenic, single-housed males. These encounters were filmed for 10 min to evaluate the behavior of the transgenic males. Behaviors such as sniffing, grooming or mounting of the female intruder were evaluated second-wise. The same female encountered two males per day, one per genotype. Between these encounters were at least 6 hours to ensure that the females did not transfer any noticeable odors from the first encounter during the second session of the day. This way we were able to compare the reaction of males of different genotypes to the same female during the same time of her estrus cycle. A total of 16 animals (8 controls/8 knockouts) were utilized.

### Female urine sniffing test

The reaction of transgenic males towards female olfactory cues was tested using female urine. For this, urine from 5–10 C57BL6J-females from different cages was collected and mixed. To circumvent reactions of the tested males towards the presentation medium, pieces of parafilm were kept in the animals’ cages for one hour prior to testing them. Then, 40 µl tap water were applied to the parafilm as neutral cue, and the time spent sniffing at the application site was assessed for 3 min. After a 45 min break, the same test was conducted using urine instead of water. A total of 16 animals (8 controls/ 8 knockouts) were utilized.

### Novelty-suppressed feeding test

Experiments were conducted according to Samuels & Hen^79^ and as visualized in Fig. 5a. In brief, mice were weighed and transferred to individual cages (IVC cages, Tecniplast) with environmental enrichments and *ad libitum* access to water. After 24 h of food deprivation, the novelty suppressed feeding test was conducted. For this, mice were individually placed in an arena (i.e., a mouse shipping container with bedding on the ground) with a food pellet attached to a platform in the center. Mice were filmed from above to later analyze their movement patterns using DeepLabCut^80^. Once a mouse started consuming the presented pellet, the animal was transferred to its home cage, and a food pellet of known weight was placed in the food tray. After the test, all animals were weighed again. In total, 20 mice (10 control/ 10 knockout) aged 3–10 months were tested.

### Luminex blood analysis

To test for hormonal blood concentrations, blood samples were collected *post-mortem*. For each mouse, two samples of 50 µl blood serum were processed. Samples were purified and analyzed using Luminex System (Millipore). Using antibody-coated beads, a multiplex assay was performed to assess changes in hormone levels being quantified using the MAGPIX detection system. The MILLIPLEX Analyst 5.1 software then calculated hormone levels of ghrelin, GLP-1, glucagon, insulin, leptin, PYY, and TNFα in pg/ml. A total of 6 control and 6 knockout mice, all aged 6 months, were used.

### Brain tissue preparation

Mice were anesthetized using 5% (v/v) isoflurane before decapitation. For immunohistochemistry, mice were transcardially perfused first with phosphate-buffered saline (PBS; pH 7.4), then with 4% paraformaldehyde (PFA) in PBS (pH 7.4). Subsequently, brains were prepared and kept in 4% PFA for 24 h at 4 °C for post-fixation. Cryoprotection was achieved by incubating the samples in 10% (v/v) and then in 30% (v/v) sucrose in PBS for 24 h each. Brains were subsequently frozen on dry ice and stored at −80 °C.

### Single-nucleus RNA sequencing

Neocortical tissue of *Pvalb-Cre/tdTomato* and *Pvalb-Cre/tdTomato/Dnmt1 loxP^2^* mice was dissected from 1-mm-thick coronal slices corresponding to the region between Bregma +1 and −2. For each genotype, tissue from the three mice was pooled prior to nuclei extraction to obtain sufficient material and minimize inter-individual variability. The samples were snap-frozen on dry ice and stored at −80 °C until nuclei extraction to preserve RNA integrity. The nuclei were extracted, fixed and stored until library preparation as previously described^12^. The snRNA-seq libraries were generated using the Evercode™ WT kit (Parse Biosciences, USA). The quality of the cDNA and the final libraries was verified using the High Sensitivity D5000 and D1000 ScreenTape kits (Agilent Technologies, USA). Libraries were equimolarly pooled and sequenced together with 5% PhiX spike-in library on an Illumina NovaSeq machine running in paired-end mode with the S2 Reagent Kit v1.5 (200 cycles; Illumina, USA).

The sequencing dataset was processed and filtered using the default parameters in Trailmaker™(https://app.trailmaker.parsebiosciences.com/; Parse Biosciences, 2025). The generated Leiden-clustered dataset was downloaded as a Seurat object and analyzed further using the Seurat^81^ package (5.3.0) in R (4.4.3). For visualization, a uniform manifold approximation and projection (UMAP) embedding was computed to represent cell clusters in a low-dimensional space. Cell clusters containing fewer than 0.3% of the total nuclei were removed to minimize potential artifacts arising from low-quality clusters. Cluster-specific marker genes were identified using the presto package implementation of the Wilcoxon rank-sum test, comparing each cluster against all other cells. The top 50 upregulated and top 50 downregulated marker genes per cluster were cross-referenced with established cell type-specific markers from published literature and publicly available reference databases.

The annotated clusters were subsequently used to infer cell–cell communication probabilities using the CellChat^82^ database and the corresponding R package (CellChat version 1.6.1).

### Immunohistochemistry

For immunohistochemistry of free-floating, 40-μm-thick, adult brain sections, slices were first washed three times in PBS/0.05% Triton X-100. GABA-staining required incubation in 1 N HCl for 10 min at 4 °C, then in 2 N HCl for 10 min at room temperature and lastly in 2 N HCl for 20 min at 37 °C. Afterwards, all sections were placed in citrate buffer (0.05% Tween20, pH 6) for 15 min at 90–100 °C. After three washing steps with PBS/0.05% Triton X-100, sections were blocked for 2 h using blocking solution (4% BSA and 10% NGS in PBS/0.05% Triton X-100). Incubation with primary antibodies was conducted for 60 h at 4 °C (rabbit anti-GABA (A2052, 1:750), rabbit anti-β-Endorphin (Ab5028, 1:1000), mouse anti-RFP (MA5-15257, 1:500)). Secondary antibodies were applied for 3 h at room temperature (Cy5-donkey anti-rabbit (A10523, 1:1000), Cy3-goat anti-mouse (115035062, 1:1000)). Sections were then stained with DAPI (100 ng/mL in PBS; Sigma-Aldrich) for 5 min. Afterwards, they were mounted onto object slides in pre-warmed 0.5% gelatine supplemented with 0.05% KCrS_2_ O_8_ x H_2_O before being embedded in Mowiol (Carl Roth).

For the staining of perineuronal nets, 30-μm-thick adult brain cryosections were collected on glass slides. Sections were washed three times in PBS/0.2% Triton X-100. After blocking for 2 h using blocking solution (4% BSA and 10% NGS in PBS/0.2% Triton X-100), the slices were incubated with Wisteria floribunda lectin (WFA; L32481, Thermo Fisher Scientific, 1:200) at 4 °C overnight. The slices were then washed three times with PBS/0.2% Triton X-100 before they were stained with DAPI (100 ng/mL in PBS, Carl Roth) for 15 min. After two washing steps with PBS, the slices were embedded in Mowiol (Sigma-Aldrich).

### Microscopy and image analysis

The brain slices were imaged using a Leica DMi8 fluorescent microscope in combination with a THUNDER® imager unit (Leica, Germany) and the corresponding software LASX (Leica, Germany). Post-processing was performed in the LASX software using the “Mosaic Merge” and “Thunder Lightning” tools. A maximum intensity projection of the merged tile scans was processed using Fiji^83^ (ImageJ, NIH, USA). Additionally, PV interneurons and surrounding PNNs were captured using a Leica SP8 Tau-STED3X confocal microscope (Leica, Germany), and the software LASX (Leica, Germany). The resulting z-stacks were processed with the Fiji software (ImageJ, NIH, USA), applying a 50-pixel sliding paraboloid background subtraction.

Fluorescence intensity measurements were performed using Fiji (ImageJ, NIH, USA). Regions of Interest (ROI) were manually defined, and the mean corrected density of each ROI was calculated by normalizing the corrected integrated density to the ROI area. For the analysis of PV interneurons enwrapped in PNNs, QuPath^84^ version 0.6.0 was used. Based on a groovy script for QuPath developed for this study, PV interneurons in the primary somatosensory cortex (S1) were identified, and the surrounding region was analyzed for the presence of WFA signals.

### Statistics

Statistical tests were carried out using Matlab (2022b), Python (2.7), and R (4.4.3). Normally distributed data was tested using two-tailed Student’s *t*-test, one-way ANOVA or two-way ANOVA. Non-normally distributed or ordinal data was tested using a Wilcoxon *rank-sum* test. Tukey’s test was conducted as a post-hoc test. The conducted tests for individual experiments are indicated in the text.

## Acknowledgments

We thank Nilufar Nojavan Lahiji and Hendro Langecker for helping with the Morris water maze experiment, Annalena Dobbert for support with the histological analysis, Mira Jakovcevski for methodical assistance in the coat state analysis, Sandra Brill for assistance in the coat state analysis, support in animal breeding/caretaking as well as fruitful scientific discussions.

## Author contributions

J.L.: conceptual design, performed experiments, data analysis, figure illustration, wrote the manuscript; C.B.Y.: conceptual design, performed experiments, data analysis, figure illustration, wrote the manuscript; K.V.: performed experiments, data analysis, figure illustration, manuscript editing; G.N.: method development, experimental supervision, data analysis, figure illustration, manuscript editing; S.G.: performed experiments, data analysis; D.P.: experimental supervision, data discussion, manuscript editing; J.R.: performed experiments, manuscript editing; M.H.: performed experiments, manuscript editing; B.K.: data discussion, manuscript editing; A.U.: performed experiments, data analysis, data discussion, manuscript editing; S.M.: performed experiments, data analysis, data discussion, manuscript editing; G.Z.B.: conceptual design, data discussion, wrote the manuscript. All authors have read and agreed to the published version of the manuscript.

## Extended Data

**Extended Data Figure 1:**
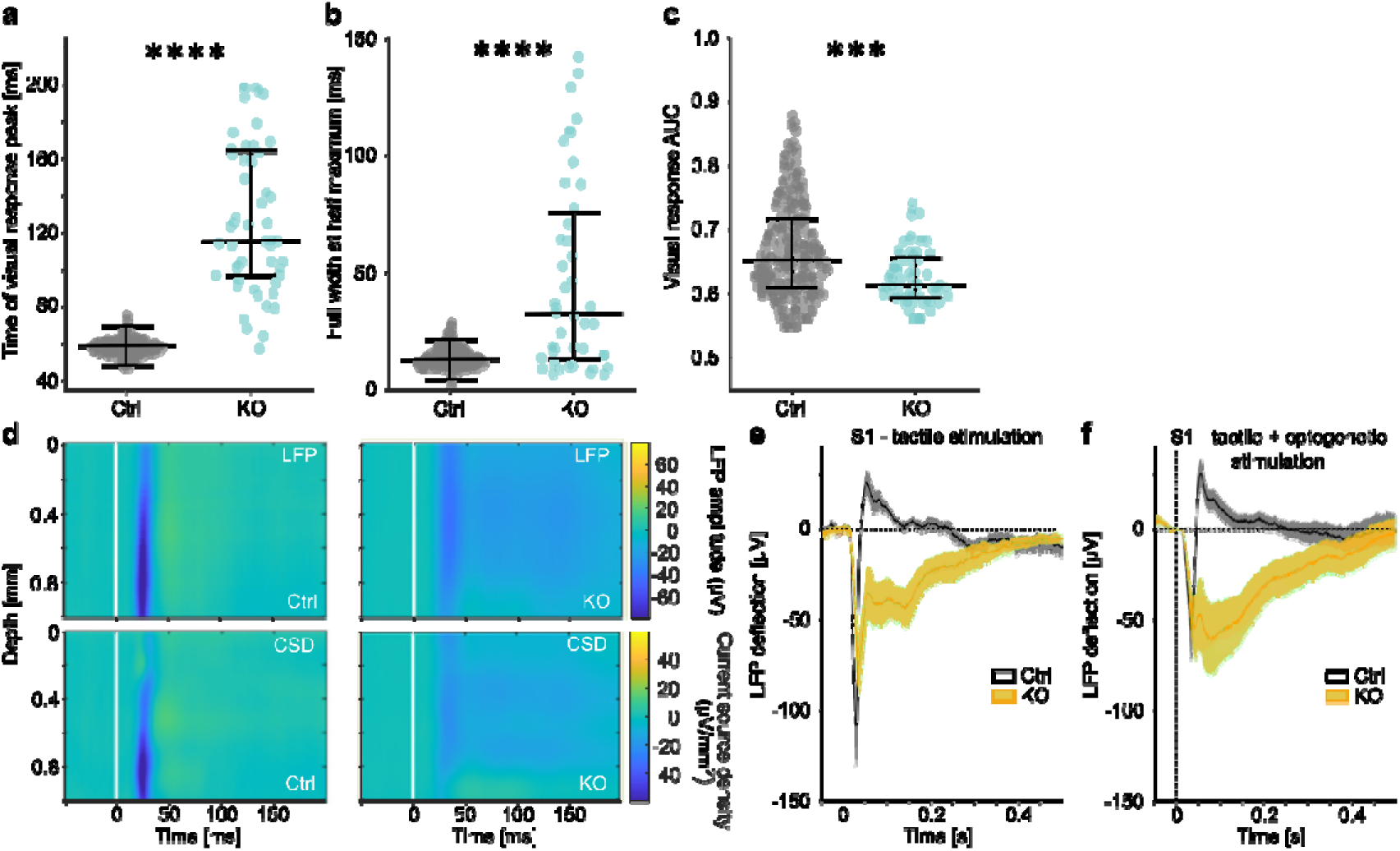
In vivo electrophysiological recordings in V1 and S1 suggest similar alterations in network processing across the cortex. Recordings were conducted in the primary visual (V1, panels a–c) or somatosensory (S1, panels d–f) cortex of awake mice (also see scheme in Fig. 1a), using either low-frequency noise patterns for visual stimulation or air puffs for tactile stimulation (*n* = 8 recordings from 2 mice per genotype). **(a–c)** Quantification of the traces shown in Figure 1e. **(a)** Time of peak firing rate in response to visual stimulation increased in *Pvalb-Cre/tdTomato/Dnmt1 loxP^2^* (KO) mice compared to *Pvalb-Cre/tdTomato* (control) mice (Wilcoxon *rank-sum* test: *p* = 2×10^-26^). **(b)** Stimulus responses of mice were less precise as reflected in an increased width at half maximum height (Wilcoxon *rank-sum* test: *p* = 1.4×10^-7^). **(c)** The area under the receiver operator characteristic curve (AUC) implies that in response to stimulation, fewer cells were involved in the cortical processing of KO mice (*n*_Ctrl_ = 31%, *n*_KO_ = 22%; Wilcoxon *rank*□*sum* test: *p* = 6.1×10^-4^) with a lower specificity (Wilcoxon *rank-sum* test: *p* = 1.4×10^-4^). **(d–f)** Local field potentials (LFP) and current source densities (CSD) of control and *Dnmt1*-KO mice recorded in S1 in response to tactile stimulation at time point 0. **(d)** LFPs and CSDs of control and KO mice across the whole depth of the cortex. White lines indicate tactile stimulus onset. **(e)** LFPs in S1 in response to tactile stimulation at time point 0, averaged across cortical layers. Shading shows SEM across recordings. **(f)** LFPs in S1 in response to tactile stimulation with concurrent optogenetic activation of PV interneurons. Panels a–c display individual data points with median (center line) and interquartile range (upper and lower lines). Ctrl = control, CSD = current source density, KO = knockout, LFP = local field potential, S1 = primary somatosensory cortex, V1 = primary visual cortex; * *p* < 0.5, ** *p* < 0.01, *** *p* < 0.001, **** *p* < 0.0001.

**Extended Data Figure 2:**
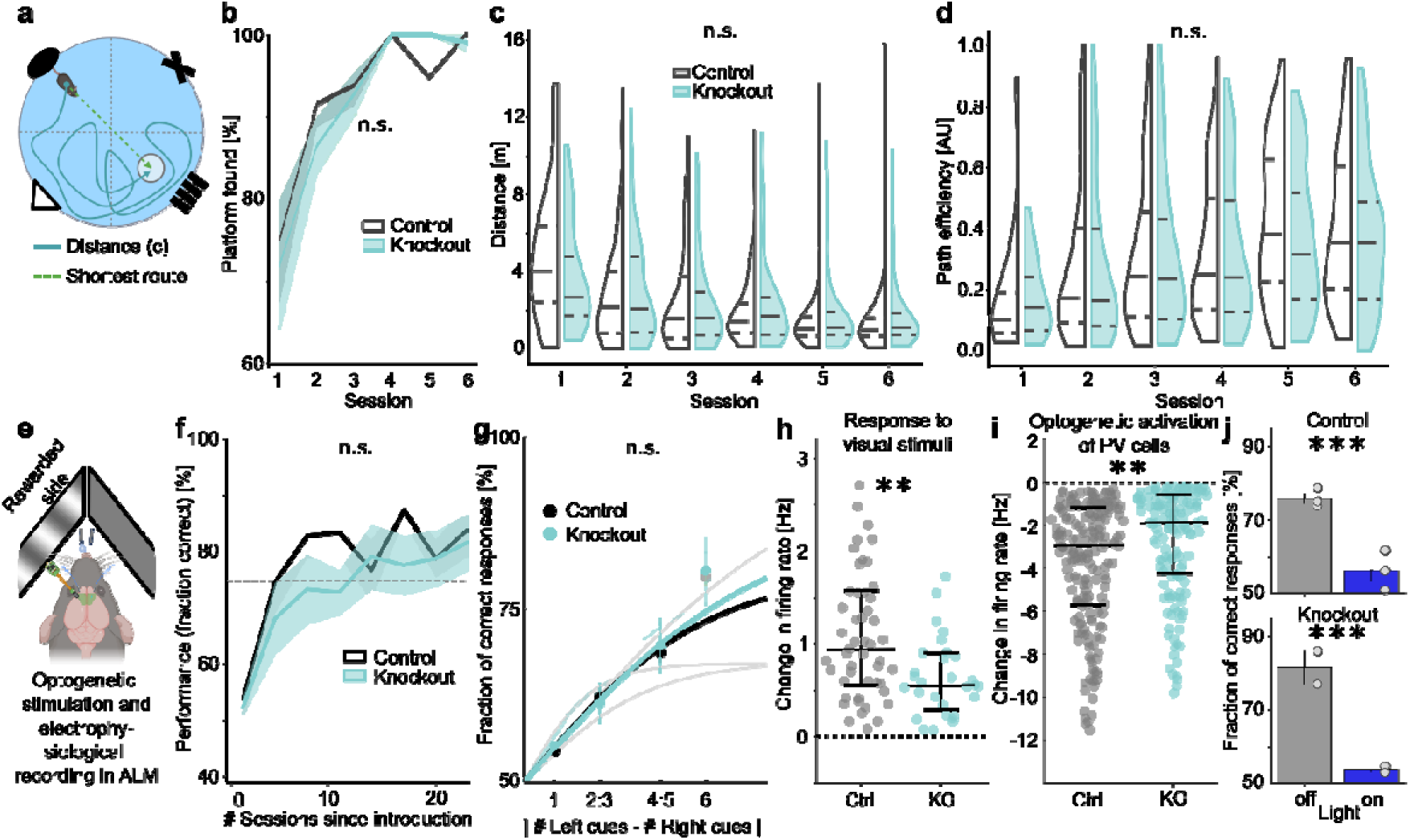
Learning and memory formation are intact in *Dnmt1*-deficient mice, also in a sensory processing-dependent task. **(a-d)** Learning and memory formation of *Pvalb-Cre/tdTomato* control (Ctrl) and *Pvalb-Cre/tdTomato/Dnmt1 loxP^2^* (KO) mice were tested in a Morris water maze (MWM), showing similar performances of both genotypes (*n* = 8 mice per genotype). **(a)** Task schematic: mice were introduced into the MWM from randomly alternating sites and had to find a hidden platform based on visual landmarks. **(b)** Mice of both genotypes similarly improved their success rate across sessions (mean ± SEM of mice; Binomial test: *p* = 0.716). **(c)** Mice of both genotypes similarly improved their swimming distances (Two-Way ANOVA (Repeated Measures): *p*_(Genotype)_ = 0.63, *p*_(Sessions)_ = 1.3×10^-22^, *p*_(Genotype x Sessions)_ = 0.92). **(d)** Path efficiencies, i.e., the swimming distance per trial divided by the respective shortest route to the platform (visualized in panel a) did not differ in knockout mice (Two-Way ANOVA (Repeated Measures): *p*_(Genotype)_ = 0.11, *p*_(Sessions)_ = 7.9×10^-15^, *p*_(Genotype x Sessions)_ = 0.36). **(e–j)** Control and KO mice were trained on a visual evidence accumulation task (*n*_(Ctrl)_ = 3, *n*_(KO)_ = 4). **(e)** Task schematic: head-fixed mice were trained on reporting the side where the higher amount of stimuli was presented on a screen by licking a corresponding water nozzle. In expert mice, electrophysiological recordings using Neuropixels probes were conducted in the anterolateral motor cortex (ALM) during multiple sessions. **(f)** Learning curves for a visual evidence accumulation task and **(g)** corresponding psychometric curves of individual animals showed no genotype-dependent effects. **(h–j)** Electrophysiological recordings in ALM of mice trained on a visual evidence accumulation task showed similar changes in cortical activity as in V1. **(h)** Firing rates of ALM neurons in response to visual stimulation in trained mice showed weaker signals in knockout compared to control mice. **(i)** Optogenetic activation of PV cells in the ALM likewise impaired decision-making in control (*n* = 3, top) and knockout (*n* = 2, bottom) mice. **(j)** Baseline-corrected firing rate of all PFC neurons that were inhibited during optogenetic activation of PV interneurons. PV-induced inhibition was significantly weaker in knockout mice compared to control animals. Panels h–i display individual data points with median (center line) and interquartile range (upper and lower lines). * *p* < 0.5, ** *p* < 0.01, *** *p* < 0.001, **** *p* < 0.0001.

**Extended Data Figure 3:**
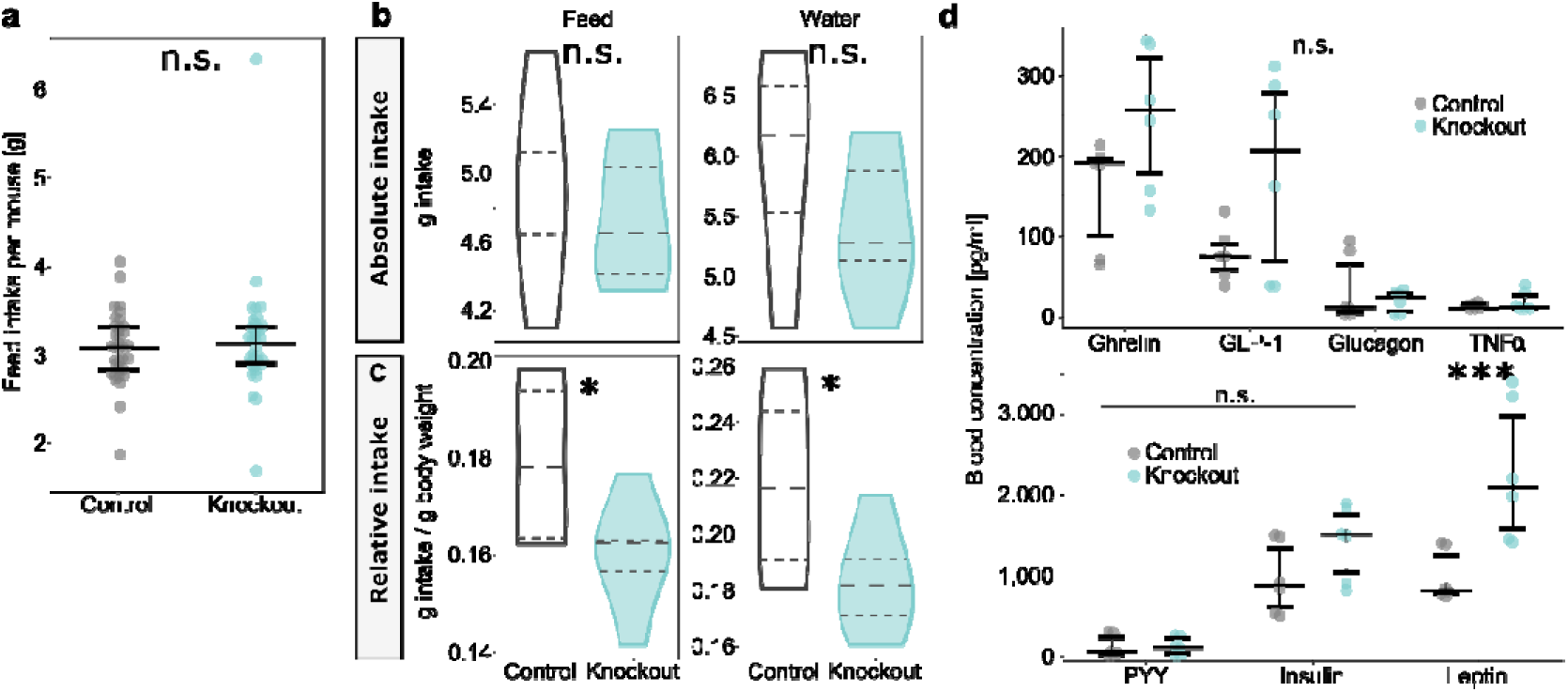
The excess body weight of *Pvalb-Cre/Dnmt1*-knockout mice is likely not appetite-based. **(a-c)** Group-housed *Pvalb-Cre/tdTomato* (control) and *Pvalb-Cre/tdTomato/Dnmt1 loxP^2^* (knockout) mice had the same food consumption rates (*n* = 8 mice per genotype). **(b)** In single-housed males (*n*_(Control)_ = 4, *n*_(Knockout)_ = 6) both the absolute feed intake as well as the absolute water intake were not altered in *Pvalb-Cre/Dnmt1*-KO mice (upper panels). **(c)** In contrast, the respective intake relative to the animals’ body weight showed a significant reduction in food and water intake in knockout mice. **(d)** Blood levels of various appetite-regulating hormones were unaltered by the *Dnmt1*-knockout (*n*_(Control)_ = 19, *n*_(Knockout)_ = 16). Prominently, only leptin levels were increased in knockout mice. Panels a and d display individual data points with median (center line) and interquartile range (upper and lower lines). * *p* < 0.5, ** *p* < 0.01, *** *p* < 0.001, **** *p* < 0.0001.

**Extended Data Figure 4:**
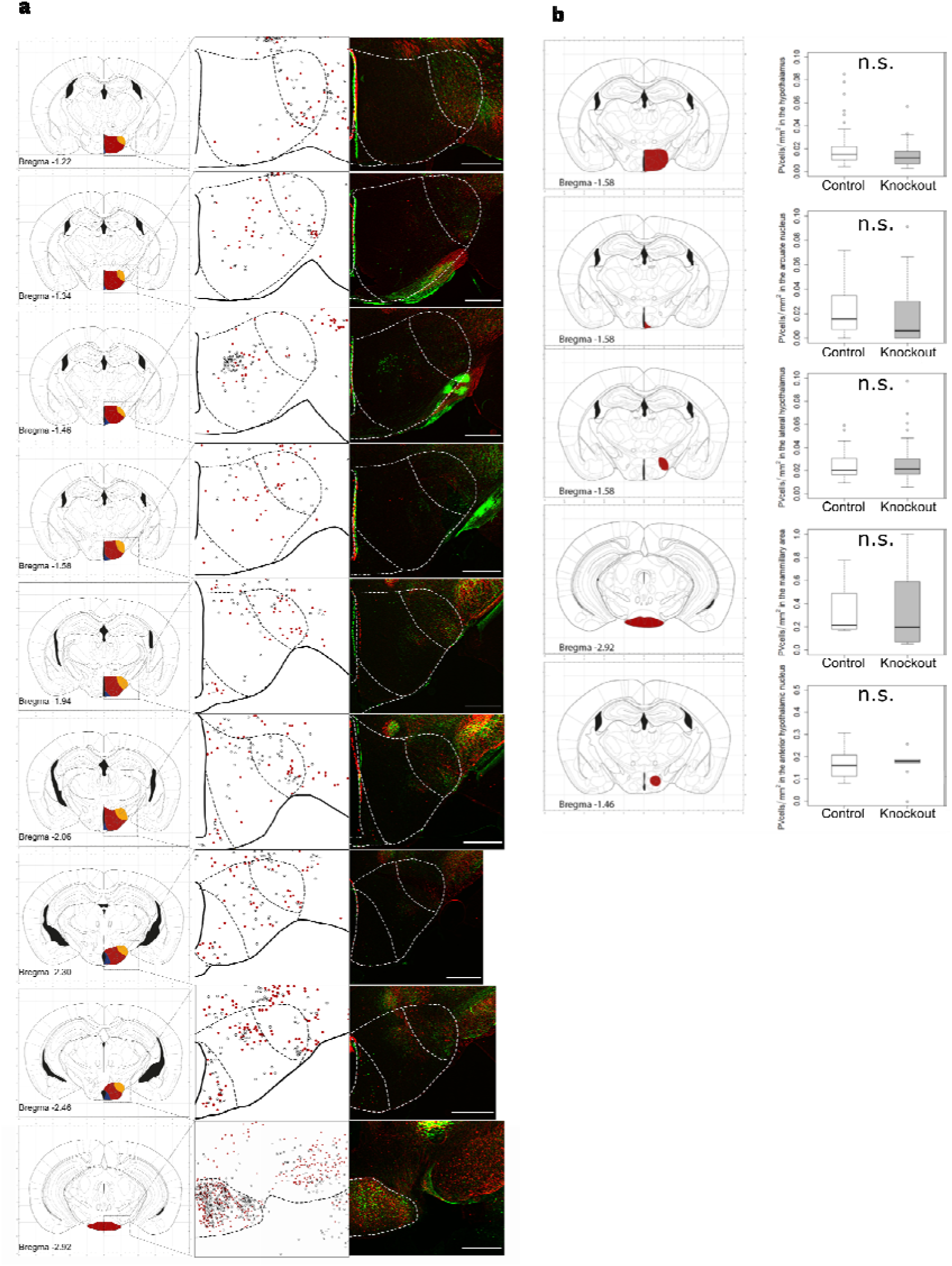
PV cell distribution and quantity were not altered in *Pvalb-Cre/tdTomato/Dnmt1 loxP^2^* (knockout) mice. **(a)** Cell distribution of tdTomato-labeled PV interneurons was unaffected by the *Dnmt1* knockout. Each row shows a subcortical region relevant in appetite regulation. Left: Schemes visualizing the investigated brain areas. Middle: Overlay of PV cell distribution found in *Pvalb-Cre/tdTomato* (control; grey dots) and knockout (red dots) mice. Right: Pseudocolor overlay of microscopy images of the investigated area. The green color indicates tdTomato-labeling in control mice, red color denotes tdTomato-positive cells in knockout mice. Scale bars = 500 µm. **(b)** Red marked regions in the illustrations on the left-hand side show areas of interest for appetite regulation, where PV cells were counted in coronal slices from different animals. Quantitative comparisons of PV interneurons in these regions revealed no differences between control and knockout animals (graphs on the right-hand side) (box plots: median, interquartile range, Tukey; *n*_(Control)_=4, *n*_(Knockout)_=3).

**Extended Data Figure 5:**
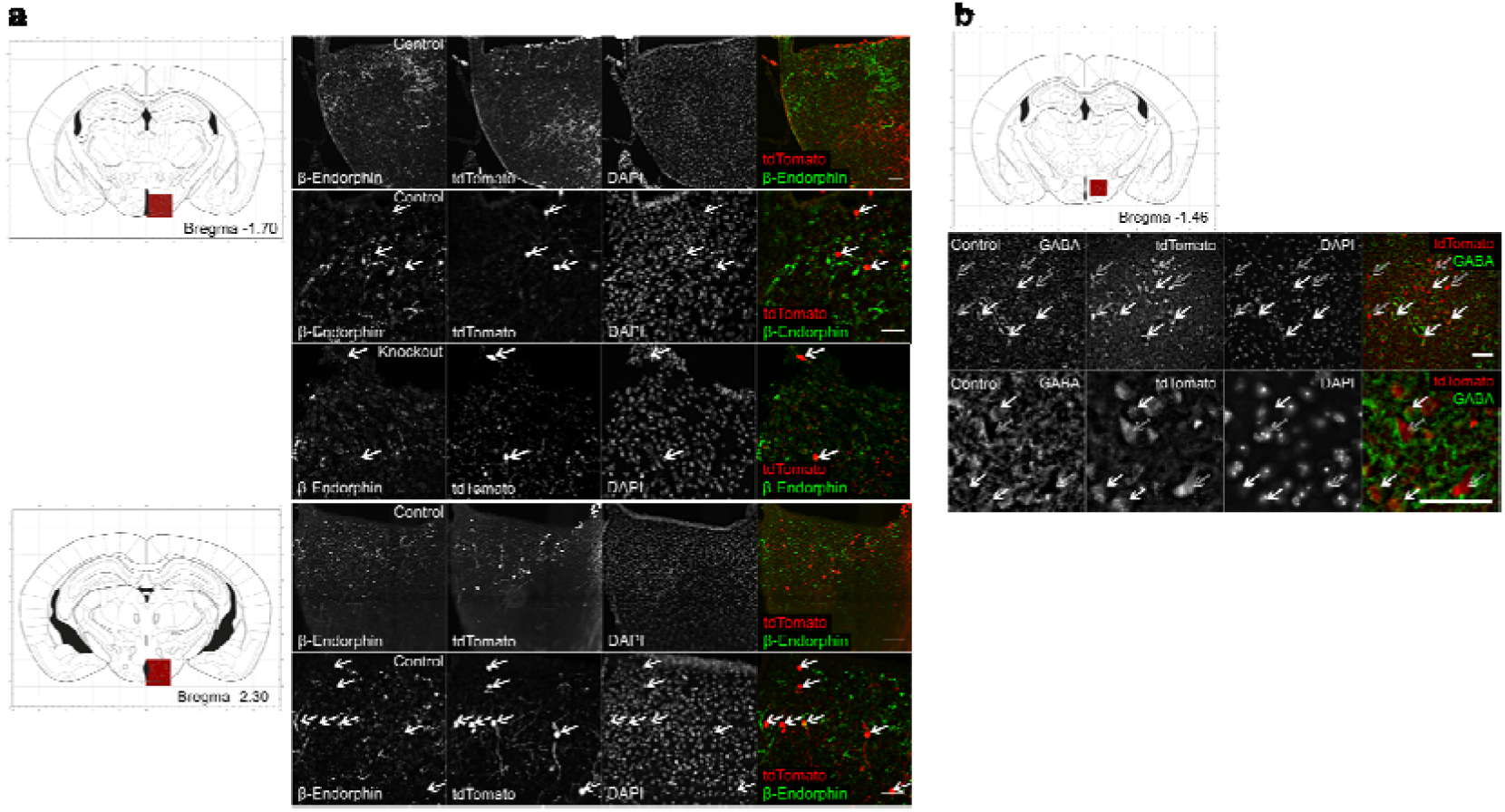
PV cell identity in the hypothalamus is unaltered in *Pvalb-Cre/tdTomato/Dnmt1 loxP^2^* (knockout) mice and does not indicate a connection between PV interneuron activity and appetite regulation. **(a)** β-Endorphin immunostaining in the arcuate nucleus of the hypothalamus was conducted to identify Proopiomelanocortin-expressing (POMC) cells. The tdTomato-labeling (white arrows) did not overlap with the β-endorphin staining in both *Pvalb-Cre/tdTomato* (control) and knockout mice (scale bar = 50 µm). **(b)** GABA immunostaining in the anterior hypothalamic nucleus of a control mouse showed a partial overlap of tdTomato- and GABA-labeling (white arrows). Immunostainings indicated that only a fraction of the tdTomato-positive PV interneurons in the hypothalamus express GABA and could thus be attributed to the group of AgRP-expressing cells, while other PV cells were not positive for GABA (empty arrows; scale bar = 50 µm).

**Extended Data Figure 6:**
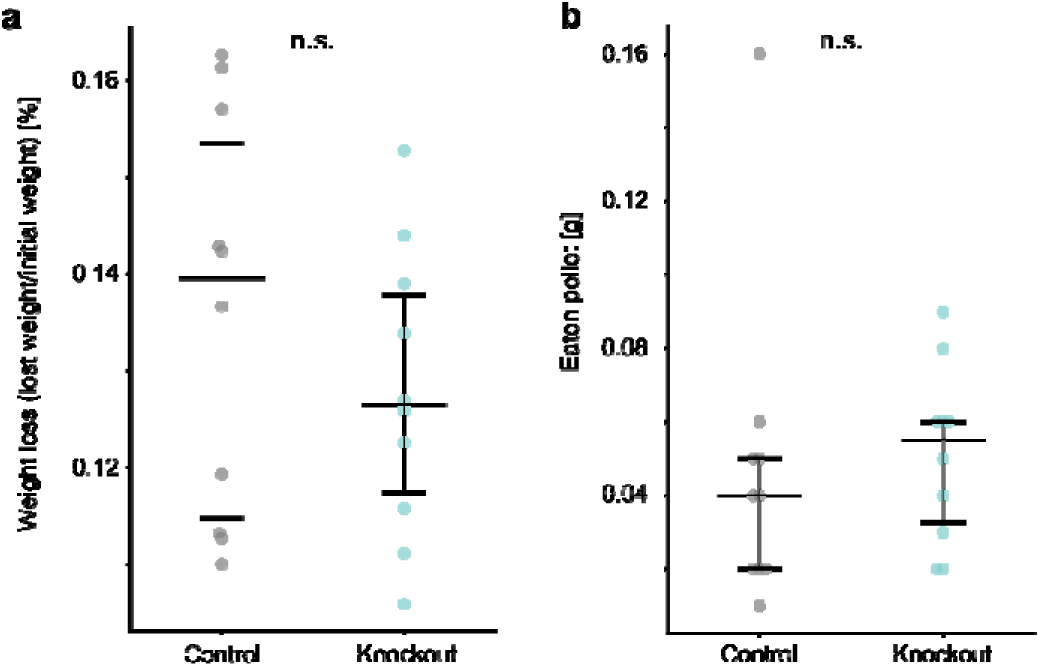
Food deprivation in the novelty-suppressed feeding test affected mice of both genotypes similarly. **(a)** *Pvalb-Cre/tdTomato/Dnmt1 loxP^2^*(knockout) and *Pvalb-Cre/tdTomato* (control) mice (*n* = 10 mice per genotype) lost similar amounts of weight after food deprivation. **(b)** In their home cages, mice of both genotypes consumed similar amounts of the pellet in a 5-min time window. Panels display individual data points with median (center line) and interquartile range (upper and lower lines).

**Extended Data Figure 7:**
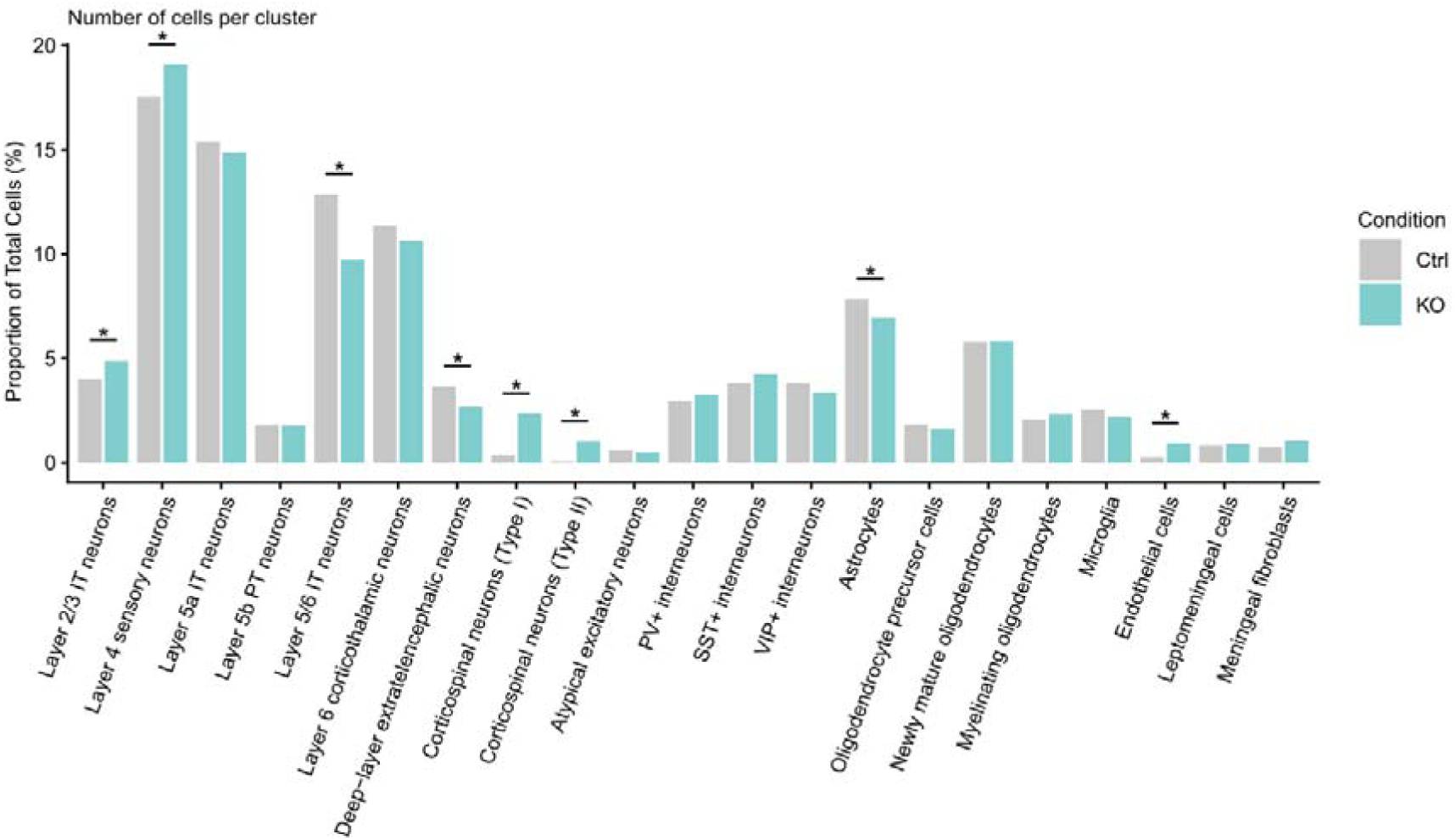
Proportions of identified clusters within the neocortex of *Pvalb-Cre/tdTomato* (Ctrl) and *Pvalb-Cre/tdTomato/Dnmt1 loxP^2^* (KO) mice normalized to the total number of cells within the given genotype. Among neuronal and glial clusters, changes were observed only for certain excitatory neuron subtypes and astrocytes (two-sample proportion test; p_L2/3 IT_ = 0.0082, p_L4_ = 0.0142, p_L5/6 IT_ = 7.0582E-12, p_Deep⍰layer ET_ = 0.0001, p_Corticospinal Type I_ = 1.8147E-30, p_Corticospinal Type II_ = 7.7081E-17, p_Astrocytes_ = 0.0417, p_Endothelial_ = 2.0998E-08,). * *p* < 0.05.

**Extended Data Figure 8:**
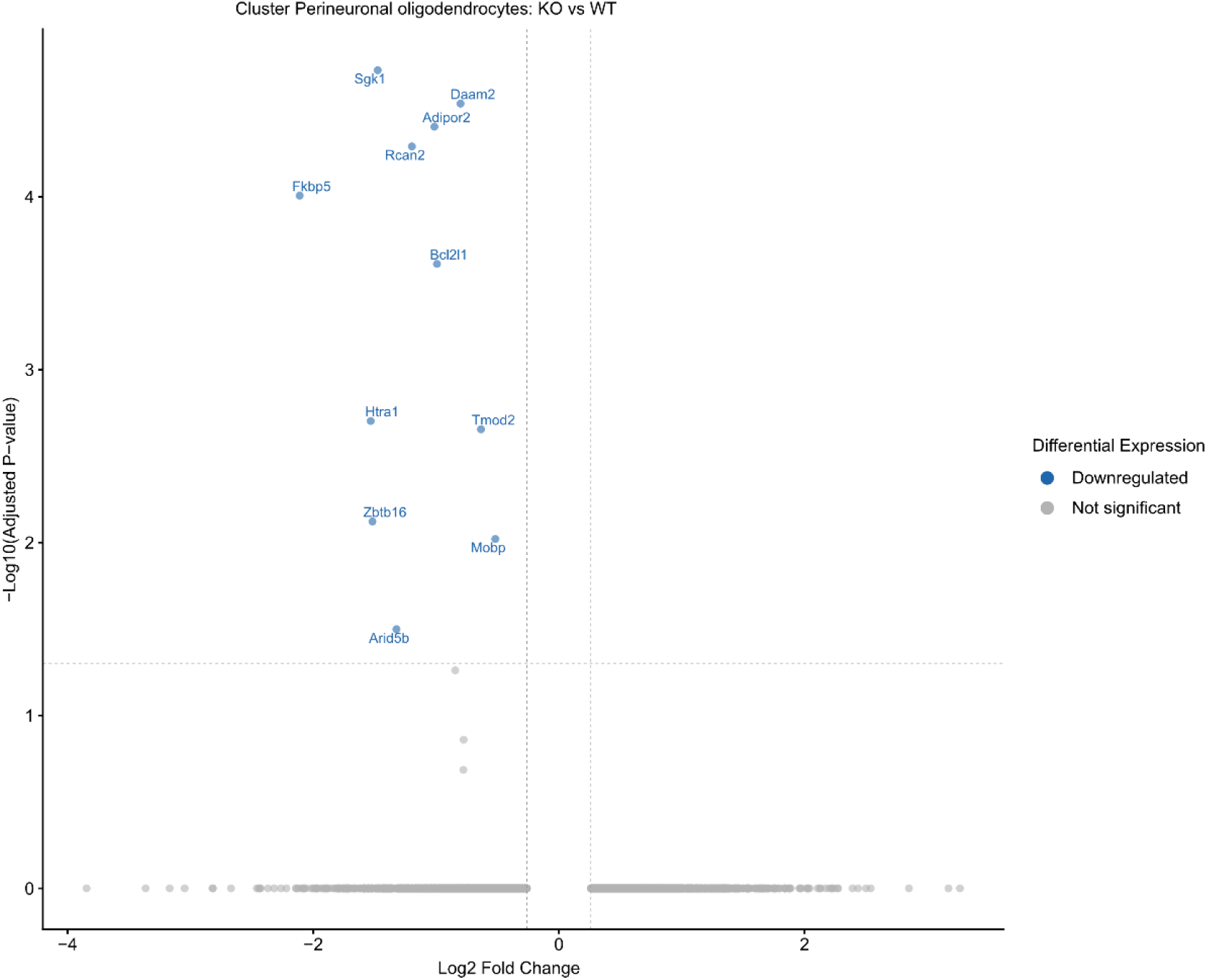
Volcano plot depicting the differentially expressed genes in the perineuronal oligodendrocytes of *Dnmt1*-KO mice. Genes implicated in stress⍰resilience, ECM⍰remodeling, and structural programs were downregulated (log_2_FC ≥ 0.26, adjusted *p* value < 0.05).

**Extended Data Table 1:** *Dnmt1*-knockout in PV interneurons induces an upregulation of genes also affected in other mouse models. Mouse lines marked with an asterisk have been found to express behavioral alterations associated with depression and/or anxiety.

| Mouse line | Enrichment<br>FDR | Number of<br>Genes | Fold<br>Enrichment |
| --- | --- | --- | --- |
| ***Htt* Mutation (HTT exon1 142Q) mouse\*** | **2.70E-06** | **21** | **5** |
| *Pmp22* KO mouse | 9.90E-05 | 16 | 4.9 |
| *Pmp22* OE mouse | 3.20E-04 | 15 | 4.7 |
| *Pmp22* KO mouse | 1.00E-03 | 13 | 4.3 |
| *Ache* AChE-S mutation mouse | 3.30E-04 | 16 | 4.3 |
| ***Htt* Mutation (HTT exon1 18Q) mouse\*** | **9.00E-04** | **14** | **4.1** |
| *Dysf* Deficiency mouse | 1.50E-03 | 13 | 4.1 |
| *Nfatc3* OE mouse | 1.00E-03 | 14 | 4.1 |
| *Tlr4* mutation mouse | 6.30E-05 | 21 | 4 |
| *Emx2* Deficiency mouse | 8.90E-04 | 15 | 4 |
| ***Dmd* Deficiency mouse\*** | **3.50E-04** | **17** | **4** |
| ***Rps6ka3* knockout mouse\*** | **3.60E-04** | **17** | **3.9** |
| ***Ache* AChE-R overexpressor mutation mouse\*** | **9.00E-04** | **15** | **3.9** |
| *Sox9* OE mouse | 8.90E-04 | 16 | 3.8 |
| *Hras* mouse | 1.20E-03 | 15 | 3.8 |
| *CHRNA9* KO mouse | 8.90E-04 | 16 | 3.7 |
| *Atxn7* OE mouse | 3.30E-04 | 19 | 3.7 |
| *Prmt5* KO mouse | 1.60E-03 | 15 | 3.6 |
| ***Sod1* G93A mutation mouse\*** | **8.90E-04** | **19** | **3.3** |
| ***Dicer1* KO mouse\*** | **1.90E-03** | **17** | **3.2** |

**Extended Data Table 2.**
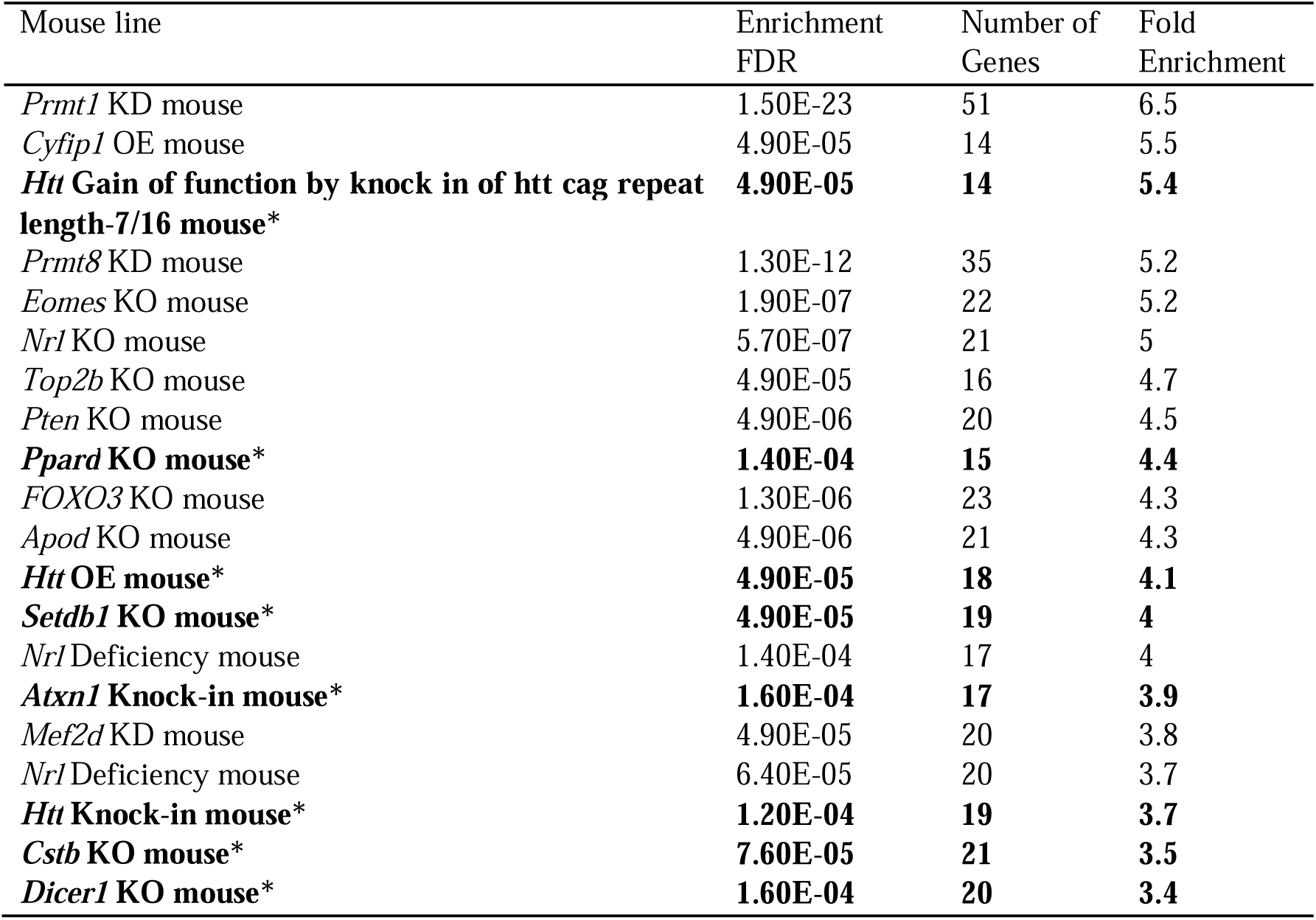
*Dnmt1*-knockout in PV interneurons induces a downregulation of genes also affected in other mouse models. Mouse lines marked with an asterisk have been found to express behavioral alterations associated with depression and/or anxiety.

